# A generalizable system for antigenic peptide targeting across HLA-I allotypes

**DOI:** 10.64898/2026.05.21.726655

**Authors:** Wyatt Blackson, Ean L. Small, Shirley M. Sun, Omkar Shinde, Ram Pantula, Sean J. Wang, Photis Rotsides, Haotian Du, Yi Sun, Daniel Hwang, Chloe S. Wang, Tianyu Lu, Eli Laskawy, Richa Kapoor, Muzamil Y. Want, Carl H. June, Regina M. Young, John M. Maris, Po-Ssu Huang, Nikolaos G. Sgourakis

## Abstract

T cell receptors (TCRs) and TCR-mimicking antibodies recognize peptide antigens in the context of specific Human Leucocyte Antigen (HLA-I) allotypes, and the extreme polymorphism of the HLA locus limits the breadth of immunotherapy development. Key barriers include divergent molecular surfaces on HLA proteins and differences in the peptide structure. As a result, existing modalities cannot confer therapeutic coverage across patients of divergent genetic backgrounds. Here, we develop an approach which combines a peptide conformational prediction tool, PepPred, with a cross-HLA binding protein engineering system, TRACeR-I^1^, to outline a generalized framework for developing binders (xTRACeRs) with compatibility across HLA allotypes while maintaining high levels of specificity towards the peptide antigen. We use our system to develop and validate xTRACeRs against clinically relevant, established peptide antigens presented across common alleles within five HLA-A/B/C supertypes^2^. Cryo-EM structures of xTRACeR-pHLA complexes for an oncofetal antigen from *PRAME* and a neuroblastoma-specific peptide from *PHOX2B* reveal effective mechanisms to navigate polymorphic HLA surface residues, and extensive interactions with the peptide. We implemented these two xTRACeRs as Chimeric Antigen Receptor (CAR) T cells and demonstrated their potent killing efficacy and specificity. Overcoming restriction across HLA supertypes lifts a key barrier in HLA-targeted immunotherapy by expanding patient coverage.

## Introduction

The cell-surface display of epitopic peptides derived from aberrant intracellular oncoproteins by HLA-I proteins^3^ forms the cornerstone of cancer immunosurveillance by T cells^4^. Substantial efforts have been invested toward developing T cell receptors (TCRs), TCR-mimicking antibodies and other modalities (reviewed in^5^) with specificity for peptide/HLA-I targets expressed in cancer and infectious disease. However, peptide target recognition through these modalities exhibits strong restriction to specific HLA allotypes^6^. HLA-I proteins are highly polymorphic, with over 20,000 allotypes present in the human population^7^, each capable of presenting millions of peptides sampled from the self and non-self (disease-related) proteome. Specific peptide sequences have the potential for presentation across multiple HLAs and show high disease recurrence (termed public antigens), which makes them attractive targets for immunotherapy development^8^. Notwithstanding, there remains a high unmet need for platforms that can address the heterogeneity of peptide/HLA molecular surfaces^9^ to develop biologics with programmable restriction against clinically relevant peptide targets.

Development of binders which could recognize the same epitope presented across different HLA allotypes necessitates that the peptide adopts a similar conformation and relies upon a binding system which confers peptide-centric specificity while maintaining compatibility with different HLA surfaces. The supertype classification of HLAs^2^ serves as an initial step for clustering allotypes based on shared overall peptide presentation (**Fig. 1a**), providing a roadmap for peptide target prioritization and cross-HLA binder development^10^. However, different HLAs exhibit polymorphic residues surrounding the peptide-binding groove which must be accommodated by any engineered biologic that seeks to overcome HLA restriction^11^ (**Fig. 1b**). This polymorphic molecular surface is compounded by the fact that peptide antigens can adopt a range of approximately 100 discrete backbone conformations, established from our analysis of solved pHLA structures^12^ (**Fig. 1c**).

**Figure 1.**
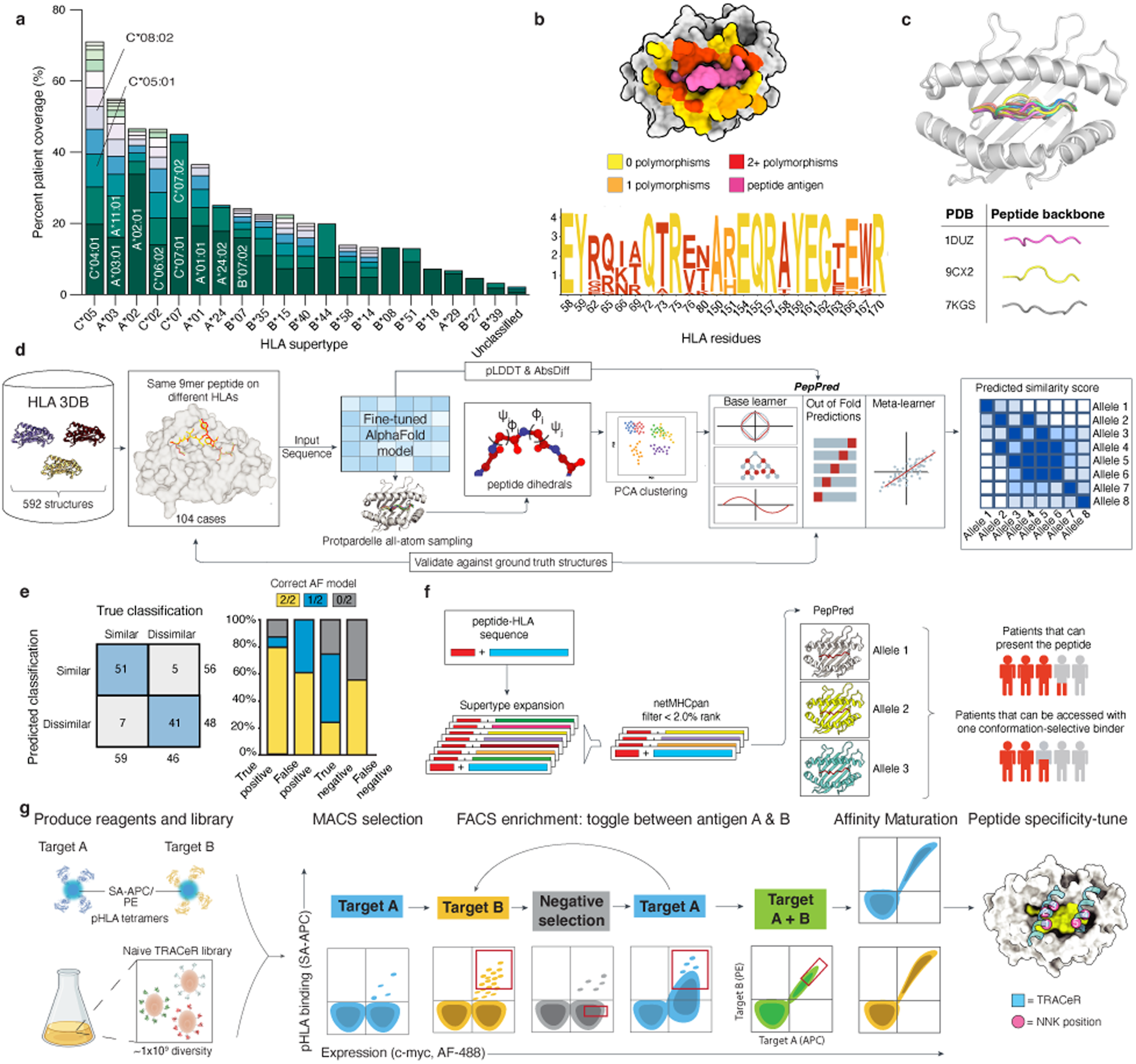
Peptide conformational prediction guides xTRACeR binder screening. **a.** Combined US population coverage for all HLA alleles which may present a similar peptide repertoire, clustered by supertype^2^. **b.** Sequence logo depicting amino acid frequencies at each HLA surface position computed across 215 common HLA-A*, HLA-B*, and HLA-C* alleles^11^. Sequence conservation across common HLA alleles is mapped onto the HLA-A*02:01 surface (PDB ID 1DUZ). **c.** Conformational diversity of peptides, highlighted in a superposition of representative structures from HLA3DB^12^. **d.** PepPred development. 9mer peptides presented on different HLAs are used to generate a ground truth set of 104 HLA pairs; their sequences are fed into AFF2^12,15,48^ to generate predicted structures and confidence metrics. Predicted structures are used to condition an all-atom generative model (Protpardelle^16^) to produce conformational ensembles via partial diffusion. Dihedral information is projected on to a low-dimensional space. Geometric features from the PCA, confidence metrics from predicted AFF2 structures, and aggregate dihedral data were used to train an ensemble model using D-Score based classification of HLA3DB structures as ground truth. A final regression meta-learner outputs all-by-all clustered heatmap based on predicted similarity scores. **e.** AFF2 performance validated against ground truth set of pHLA structures bucketed by PepPred confusion-matrix classes. **f.** Computational workflow to identify cross-HLA peptide candidates that could enable high patient coverage. Sequences of pHLAs are expanded to include other alleles in the supertype and filtered using netMHCpan^17^ binding probabilities. Sequences are fed into AFF2/Protpardelle, and PepPred to generate structure models and classify peptide conformations. **g.** Experimental workflow for generating cross-HLA binders. A yeast library containing TRACeRs with diversified amino acids at eight positions is enriched for binding to orthogonal pHLA tetramers. Specificity across alleles is selected by toggling between enrichment for each allele along with negative selections against off-target peptides. High specificity clones are affinity matured using error-prone mutagenesis.

With these barriers in mind, we leverage TRACeR-I (targeted recognition of antigen-MHC-I complex reporter), a platform that enables programmable development of pHLA-I binders using a peptide-centric docking mode and a diversified peptide specificity “box” of 8 residues^1^. We reasoned that polymorphic residues can be well accommodated by the diversified TRACeR library, as our approach has been shown to derive highly specific binders for targets spanning HLA-A*, - B*, and -C* allotypes^1^. Here, to identify cases of peptides which may adopt conserved conformations and could serve as targets for cross-HLA binder development *de novo*, we generated a structural similarity prediction tool, PepPred. Integrating TRACeR-I with PepPred, we outline a complete process for developing cross-HLA binders which retain high peptide specificity and can be leveraged as antigen binding modules for Chimeric Antigen Receptor (CAR) development. Our work establishes a generalizable avenue for expanding T cell therapy to intracellular oncoproteins across patients of diverse genetic backgrounds, directly overcoming the HLA restriction barrier.

## Results

To evaluate the extent to which peptides may adopt a similar conformation when presented on different HLA allotypes, we compared cases where the same peptide has been resolved in multiple pHLA complexes from our curated database, HLA3DB^12^ (**Extended Data Fig. 1**). Analysis of 104 such cases spanning multiple HLA allotypes shows that that 46% of pairs adopt a similar peptide backbone conformation reflected by D-scores (a metric for peptide structural divergence; lower scores indicate higher similarity^13^) of less than 1.5, as benchmarked against previously known structures^12^ (**Extended Data Fig. 1a-c**). Given this collection of structural data illustrating a high prevalence of peptides with conserved presentation across HLA allotypes (**Extended Data Fig. 1**), we developed a deep learning model to predict peptide/HLA conformational ensembles^14^ and identify relevant targets for cross-allelic binder development. The classifier algorithm, PepPred, consists of finetuned AlphaFold2^15^ (AFF2) and all-atom Protpardelle^16^ models, both trained on all 9mer pHLA structures in HLA3DB^12^ (*Appendix*). AFF2 provides initial predictions of pHLA complexes, and these structures are processed through Protpardelle via partial diffusion to sample local peptide conformations restricted by the HLA binding groove. To consider peptide flexibility in differentiating similar conformations presented on different HLA allotypes, predicted ensembles were projected onto a reduced dimensionality space computed from backbone dihedrals, and their pairwise structural similarity was evaluated using an average distance score. Gradient boosting on AFF2 confidence metrics and the predicted ensemble structural similarity, over our curated set of 104 pairs of pHLA structural cases as ground truth, was used for training. Overall, PepPred receives a pair of generated pHLA ensembles and their confidence metrics as inputs to assess the likelihood of conserved presentation between two HLA allotypes (predicted similarity score). Results computed for all HLA pairs within a supertype predicted to present a candidate peptide antigen are then plotted as heatmaps (**Fig. 1d**). PepPred was trained using multiple cross validation to diminish overfitting, resulting in an 86% true positive and 85% true negative prediction rate, despite any errors present in the initial AFF2 models (**Fig. 1e**).

Leveraging our method for identifying peptide antigens with conserved structural presentation across distinct HLA allotypes, we propose a generalizable, end-to-end workflow that enables the development of peptide-specific, cross-HLA binders. First, the overall HLA coverage of a candidate peptide target experimentally determined to be presented by any HLA allotype is predicted using netMHCpan^17^ (**Fig. 1f**). All predicted HLAs are then evaluated using PepPred to identify pHLA pairs with high likelihood for conserved peptide presentation (**Fig. 1d and f**). Fluorescent pHLA tetramers are produced recombinantly and used to enrich a yeast TRACeR library for peptide-specific cross-HLA binders (**Fig. 1g**). Initial selection is performed using magnetic activated cell sorting (MACS) and subsequent rounds of florescence activated cell sorting (FACS) are done with tetramerized pHLAs. Once clonal pools are sufficiently enriched, dual-fluorophore sorting experiments can be employed to isolate TRACeRs with a distinct cross-HLA binding pattern. High specificity binders from the initial enrichment process are then affinity matured using random mutagenesis. Further engineering through structure-guided library design can be integrated to reduce the likelihood of off-target reactivity with sequence-similar peptides (**Fig. 1g**).

To validate our approach and demonstrate its cross-HLA capability, we focused on a clinically relevant PRAME antigen (ELFSYLIEK) which shows confirmed binding to HLA-A*11:01 and HLA-A*03:01, members of the A*03 supertype (**Fig. 1a**) in a conserved conformation^18^ (**Extended Data Fig. 2**). Although an HLA-A*02:01-restricted PRAME TCR has demonstrated significant clinical efficacy in melanoma, A*03:01- and A*11:01-restricted TCRs targeting the ELFSYLIEK epitope have not been advanced to clinical development^19^. A naïve discovery effort yielded a single clone (xTR-PRAME) with desired on-target specificity (**Extended Data Fig. 3**).

To enable downstream therapeutic applications, we affinity matured this clone (xTR-PRAME^HA^) using error prone mutagenesis (**Extended Data Fig. 4**). We further observed similar binding levels to PRAME/HLA-A*68:01 tetramers, expanding therapeutic coverage to 36.8% (**Fig. 2a and b**). Beyond the known peptide similarity between HLA-A*11:01 and HLA-A*03:01, the observed HLA cross-reactivity pattern of xTR-PRAME^HA^ aligns with peptide similarity predictions made using PepPred, which suggest a main cluster that includes HLA-A*11:01/A*03:01/A*68:01 (**Extended Data Fig. 5**). Notably, HLA-A*11:01, A*03:01, and A*68:01 belong to distinct T cell seroreactivity groups^11,20^, showing that our engineered proteins can navigate polymorphic barriers through a mechanism distinct from TCR alloreactivity^18^. An alanine scan of the PRAME antigen sequence reveals targeted recognition for all peptide residues, with high specificity for positions 3, 4, 5, 6, and 7 (**Fig. 2c**). Furthermore, titration of monovalent pHLAs on yeast shows low nanomolar binding affinities, while maintaining precise cross-allotype specificity (**Fig. 2d**).

**Figure 2.**
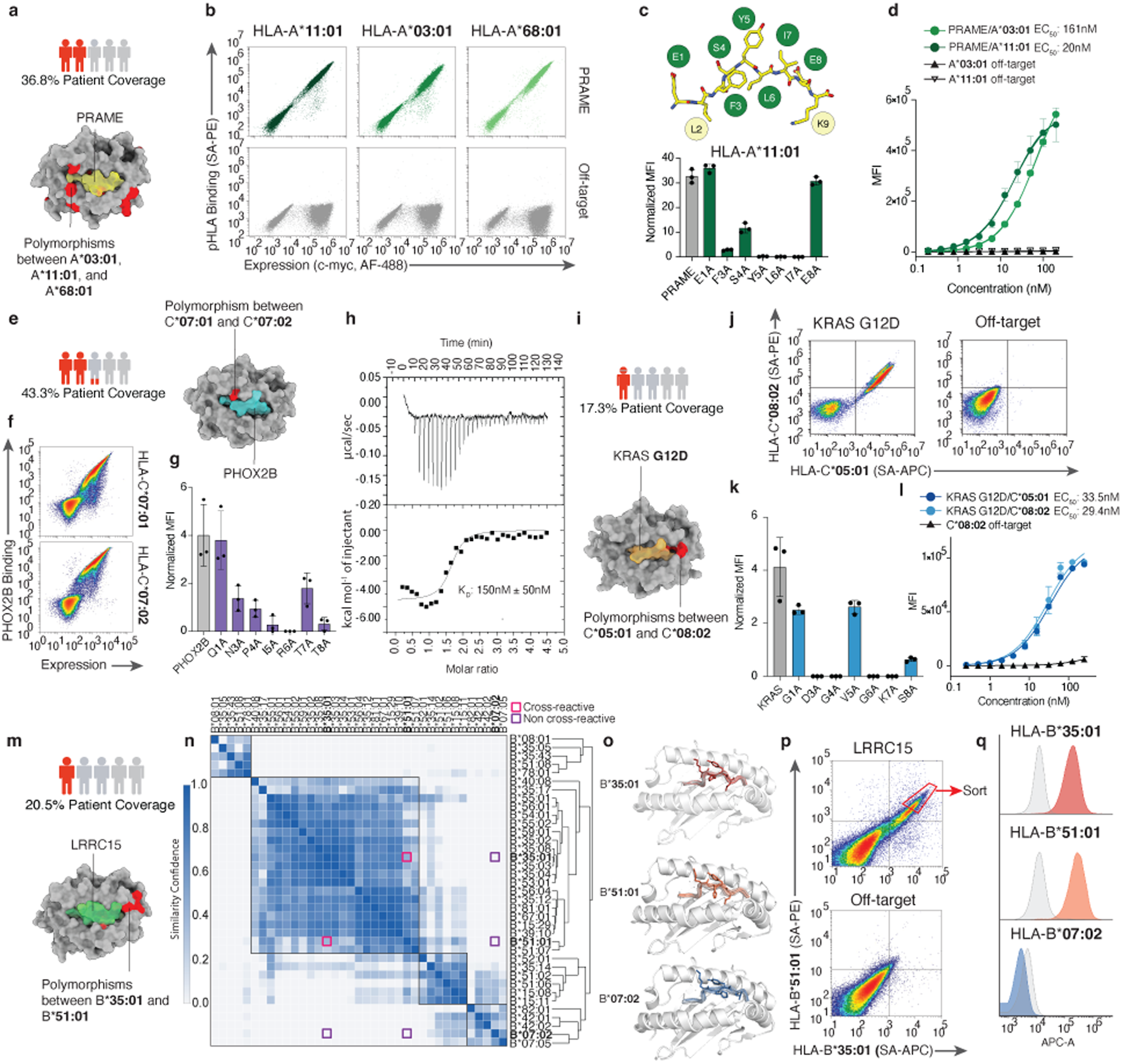
Development of xTRACeRs against clinically validated epitopes. **a.** Structure of HLA-A*03:01/PRAME showing polymorphic residues between HLA-A*03:01/A*11:01/A*68:01. **b.** Representative flow cytometry plots showing binding of xTR-PRAME^HA^ to on/off-target pHLA tetramers. **c.** Alanine scan of PRAME peptide on HLA-A*11:01 tetramers used to stain xTR-PRAME^HA^ expressed on yeast. MFI shown is mean ± SD from n=3 replicates. **d.** Yeast titration curves of xTR-PRAME^HA^ against on/off-target monovalent pHLAs. Data shown represent the mean ± SD from n=3 replicates. **e.** Model of PHOX2B/HLA-C*07:02 highlighting polymorphic residues between HLA-C*07:01/C*07:02 and total patient coverage. **f.** Representative flow cytometry data for staining of xTR-PHOX2B^HA^ with indicated pHLA tetramers on yeast. **g.** Alanine scan of PHOX2B peptide on HLA-C*07:01 tetramers used to stain xTR-PHOX2B^HA^ expressed on yeast. MFI shown is mean ± SD from n=3 replicates. **h**. Recombinant xTR-PHOX2B^HA^ exhibits nanomolar-range binding affinity via isothermal titration calorimetry (ITC). Data shown represent the mean ± SD from n=2 independent replicates. **i.** Structure of KRAS-G12D/HLA-C*08:02 highlighting polymorphic residues between HLA-C*08:02 and C*05:01 and expected patient coverage for both alleles. **j.** Representative flow cytometry plots of dual-fluorophore tetramer binding experiments. **k.** Alanine scan of KRAS-G12D peptide on HLA-C*08:02 tetramers used to stain xTR-KRAS^HA^ expressed on yeast. MFI shown is mean ± SD from n=3 replicates **l.** Yeast titration curves of xTR-KRAS-G12D^HA^ against on/off-target monovalent pHLAs. Data shown represent the mean ± SD from n=3 technical replicates. **m.** Model of LRRC15/HLA-B*07:02 highlighting polymorphic residues across HLA-B*07:02, B*35:01, and B*51:01 in red with expected patient coverage. **n.** PepPred structural similarity scores are plotted for all HLA pairs from the B07, B35, and B51 supertypes. Magenta squares indicate HLA allotypes with confirmed cross-reactivity for the same TRACeR binder, while purple squares are non-reactive HLA pairs. Hierarchical clustering of HLAs according to PepPred pairwise scores is shown on the right, and main clusters are highlighted with black boxes. **o.** Protpardelle backbone ensembles for LRRC15 on HLA-B*35:01, HLA-B*51:01, and HLA-B*07:02. **p.** LRRC15 selection via dual-fluorophore staining enables the isolation of cross-allelic TRACeRs. **q.** Representative MFI distributions for yeast staining of xTR-LRRC15 with tetramers presenting LRRC15 on indicated HLA alleles.

We then explored whether the TRACeR platform could be generalized to develop cross-allotype binders against established cancer-associated antigens presented by different HLA supertypes. The PHOX2B peptide (QYNPIRTTF) represents a *bona fide* neuroblastoma oncofetal antigen^21^. A peptide-centric CAR targeting PHOX2B can clear tumors in xenograft models, however, its therapeutic coverage is limited to carriers of HLA alleles from the A9 serological group^20^ (A*24/A*23 supertype), which represents 26% of the US population^22^. Presentation of the same epitope on C*07 supertype alleles (**Fig. 1a**), with a combined coverage of 43%, offers a path to therapy expansion (**Fig. 2e**). Binder discovery followed by affinity maturation yielded xTR-PHOX2B^HA^, a C*07 supertype cross-HLA binder against PHOX2B (**Fig. 2f; Extended Data Fig. 6**). Alanine scanning revealed varying levels of contact dependence across positions 3 to 8 of the peptide sequence, with an essential role for R6 (**Fig. 2g**). In alignment with our PepPred modeling showing a discrete clustering of HLA-C*07:01 and HLA-C*07:02, xTR-PHOX2B^HA^ can cross-react with both C*07 but not with A9 group HLAs (**Fig. 2f; Extended Data Fig. 7**). When purified recombinantly, xTR-PHOX2B^HA^ exhibited a monovalent binding affinity of 150 nM by isothermal titration calorimetry, which is suitable for CAR-T development^23^ (**Fig. 2h; Extended Data Table 1**).

The KRAS-G12D neoantigen (GADGVGKSAL) has confirmed presentation on both HLA-C*08:02 and HLA-C*05:01^24^, with existing crystallography data showing that both alleles can sample a common peptide structure despite the presence of conformational flexibility^25,26^ (**Fig. 2i; Extended Data Fig. 8**). Individual TCRs recognize KRAS-G12D restricted to either allotype, a selectivity governed by the polymorphic residues at positions 77 and 88 on the HLA surface^27^ (**Fig. 2i**). Enrichment against both allotypes presenting KRAS-G12D yielded several clones with peptide specificity (**Extended Data Fig. 9**). Affinity maturation of the most prevalent clone (xTR-KRAS) yielded several binders with increased affinity while maintaining on-target specificity (**Fig. 2j; Extended Data Fig. 10**). The lead candidate, xTR-KRAS^HA^, showed specific cross-HLA binding via a dual-fluorophore experiment in which yeast expressing xTR-KRAS^HA^ were incubated with a 1:1 ratio of orthogonal pHLA tetramers conjugated to either R-phycoerythrin (PE) or allophycocyanin (APC) (**Fig. 2j**). Peptide alanine-scan experiments demonstrated that xTR-KRAS^HA^ has specificity for surface-exposed peptide residues except position 5 (**Fig. 2k**). On-yeast titrations illustrated low nanomolar on-target binding across both HLAs presenting the KRAS-G12D neoantigen while discriminating against the KRAS-WT self-peptide (**Fig. 2l**).

Finally, we sought to implement our end-to-end workflow for the development of cross-HLA TRACeRs *de novo*, for pHLAs of unknown structures. Leucine-rich repeat-containing protein 15 (LRRC15) is a promising immunotherapy target due to its overexpression in multiple mesenchymal-derived tumor types^28^. An epitopic peptide from LRRC15 (MPLKHYLLL) was previously identified from monoallelic HLA-B*51:01 HeLa cells^29^. *In vitro* refolding assays and thermal stability measurements confirm the formation of stable protein complexes for common alleles HLA-B*07:02, HLA-B*35:01, in addition to HLA-B*51:01 (**Extended Data Table 2**). We employed PepPred to evaluate the likelihood for conserved presentation of LRRC15 across different HLAs from the three supertypes. A heat plot of peptide similarity scores reveals four well-defined HLA clusters, separating HLA-B*35:01 and HLA-B*51:01 from HLA-B*07:02, with the main B*35/B*51 cluster combining for 20.5% population coverage (**Fig. 2m and n**). This is also reflected in Protpardelle conformational ensembles, showing divergence in backbone conformations for solvent exposed peptide residues between the three alleles (**Fig. 2o**). A TRACeR discovery campaign using both single-color and dual-fluorophore sorting yielded a single clone, xTR-LRRC15 with on-target specificity for both HLA-B*35:01/LRRC15 and HLA-B*51:01/LRRC15, and no binding to HLA-B*07:02/LRRC15 (**Fig. 2p and q; Extended Data Fig. 11**), in agreement with our PepPred results (**Fig. 2n and o**). Notably, orthogonal TRACeR binders against LRRC15/HLA-B*07:02 could be isolated on yeast, further supporting that the lack of cross-HLA reactivity is due to changes in the peptide conformation, rather than a limitation of the TRACeR-I library (**Extended Data Fig. 12**). Together, our results across four clinically relevant targets indicate that our workflow can be used to streamline the development of specific binders targeting a common peptide conformation presented on HLAs across a given supertype.

To elucidate the molecular basis of cross-HLA PRAME antigen recognition by xTR-PRAME^HA^, we determined the structure of xTR-PRAME^HA^/PRAME/HLA-A*11:01/β_2_m complex using cryo-EM at 3.96 Å resolution (**Extended Data Fig. 13 and 14**). The xTR-PRAME^HA^ comprises a tandem pair of three α-helical bundles that dock perpendicularly to the α_1_ and α_2_ helices of its pHLA target using a peptide-centric binding mechanism (**Fig. 3a**). The N133 sidechain from xTR-PRAME^HA^ forms a hydrogen bond with the backbone carbonyl group from S4, while the TRACeR helices form a hydrophobic pocket around the bulged ^5^Y-L-^7^I peptide motif. We also observe R160 and R167 of the TRACeR forming salt bridges with E8 of PRAME (**Fig. 3b**). The intolerance to Ala substitution at P3 (**Fig. 2c**), despite the absence of direct contacts, suggests that Phe at this position allows the peptide to adopt a conformation compatible with xTR-PRAME^HA^ recognition. Together, our structural findings reveal that TRACeR leverages multiple interactions and shape complementarity with the peptide, and these features suggest high antigen specificity. To define the structural basis for multi-allotype compatibility, we highlighted polymorphic residues among HLA-A*03:01, HLA-A*11:01, and HLA-A*68:01 on the complex structure. We observed that xTR-PRAME^HA^ engages exclusively conserved residues on the α_1_ helix, by leveraging a helical motif which crosses the HLA groove between the main polymorphic clusters on both ends of the α_1_/α_2_ helices (**Fig. 3c**).

**Figure 3.**
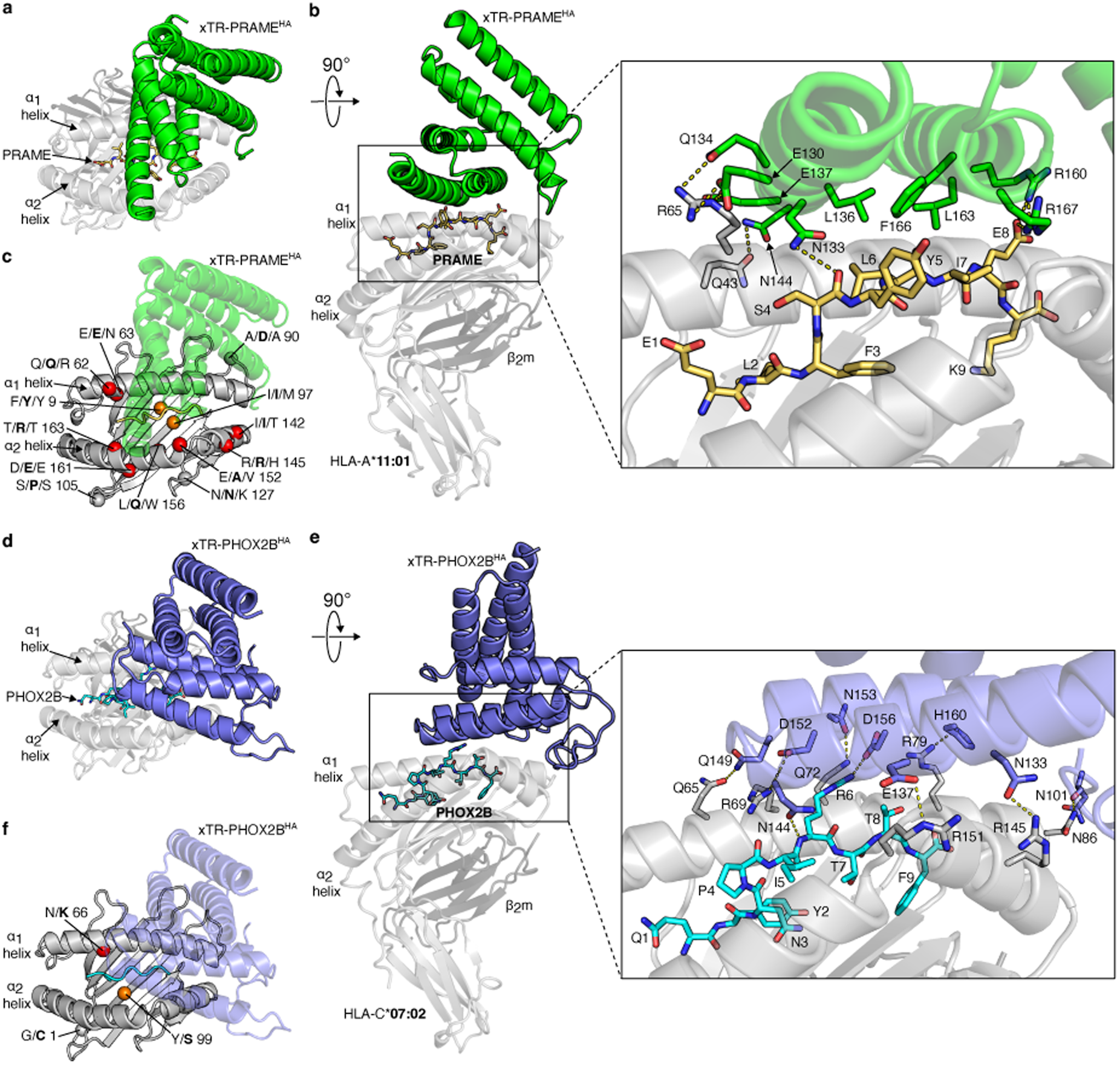
Structural basis for PRAME and PHOX2B antigen recognition by xTRACeRs. **a.** Top-view of xTR-PRAME^HA^ bound to PRAME/HLA-A*11:01/β_2_m. The PRAME peptide is shown in yellow. **b.** Side-view of the complex in panel a showing that the xTR-PRAME^HA^ adopts a peptide-centric docking orientation. Inset showing atomic interactions formed between the xTR-PRAME^HA^ sidechains with key peptide and HLA residues. Yellow dashed lines represent hydrogen bonds. **c.** HLA polymorphisms among the HLA-A*03:01, HLA-A*11:01, and HLA-A*68:01 allotypes are shown as spheres and colored according to their position on the HLA surface. (Red: α_1_ and α_2_ helices; Orange: groove; Grey: connecting loop). HLA-A*11:01 residues are shown in bold. **d.** Top-view of xTR-PHOX2B^HA^ bound to PHOX2B/HLA-C*07:02/ β_2_m. PHOX2B epitope is shown in cyan. **e.** Side-view of the complex in panel d showing peptide-centric docking orientation. Inset shows the interactions mediated by xTR-PHOX2B^HA^ with the key peptide and HLA residues. **f.** HLA polymorphisms present in HLA-C*07:01 and HLA-C*07:02 are shown as spheres and colored according to panel their position on the HLA surface as in panel c. HLA-C*07:02 residues are shown in bold.

We further determined the cryo-EM structure of xTR-PHOX2B^HA^/PHOX2B/HLA-C*07:02/β_2_m complex at 3.89 Å resolution (**Extended Data Fig. 15 and 16**). The structure reveals that the overall domain architecture of the TRACeR scaffold is maintained relative to both the xTR-PRAME^HA^ and our previous X-ray structure of a prototypical domain-swapped TRACeR bound to NY-ESO-1/HLA-A*02:01^1^ (**Extended Data Fig. 17a**). Notably, the xTR-PHOX2B^HA^ undergoes a rigid-body rotation to adopt a docking mode which places the two TRACeR helices in alignment with the peptide-binding cleft of PHOX2B/HLA-C*07:02 (**Extended Data Fig, 17c**). This stacked-helix arrangement promotes an optimized peptide interface, while avoiding direct contacts with the dimorphic HLA residue at position 66 (**Fig. 3d-f**). Peptide-specific recognition, supported by our peptide Ala scan experiments (**Fig. 2g**), is conferred by a salt bridge between D156 and R6 from PHOX2B and further supported by hydrogen bonds and hydrophobic stacking interactions between F140, N144 and the bulged peptide backbone, in an analogous manner to PRAME recognition by xTR-PRAME^HA^ (**Fig. 3e; Extended Data Fig. 18**). We further observe that docking of xTR-PHOX2Bᴴᴬ is mediated through extensive contacts with HLA-I residues that are conserved between HLA-C*07:01 and HLA-C*07:02 (**Fig. 3f; Extended Data Fig. 18b**). Structural comparison between our xTR-PHOX2Bᴴᴬ/HLA-C*07:02 complex and our previous co-crystal structure of the 10LH PC-CAR ScFv/HLA-A*24:02 complex^22^ reveals divergent peptide conformations (D-score 5.97), in alignment with our PepPred predictions (**Extended Data Fig. 7 and 19**). This result further highlights that barriers to target recognition across different HLA allotypes are imposed by changes in presentation of the peptide (**Extended Data Fig. 19**).

To complement and further validate our structural studies, we characterized the peptide specificity profile of xTR-PHOX2Bᴴᴬ *in vitro*. First, we generated a synthetic peptide library comprising all single amino acid substitutions of non-anchor residues in the PHOX2B sequence^22^. Pulsing of each peptide from our library on HLA-C*07:02 monoallelic B721.221 cells^30^, followed by staining with tetramerized xTR-PHOX2B^HA^, revealed high binding specificity for residues 4, 5, 6, and 8 (**Extended Data Fig. 20)**, in agreement with the docking mode and contacts identified in our cryo-EM structure of the cognate complex and our Ala scanning experiments (**Fig. 3e and 2g**). In contrast, binding of xTR-PHOX2B^HA^ was tolerant to specific amino acid substitutions at residues 1, 3, and 7, that are oriented towards the peptide-binding groove (**Fig. 3e; Extended Data Fig. 20 and 18b**).

Following incorporation into a second generation 4-1BB/CD3ζ CAR^31^ construct (**Fig. 4a**), xTR-PHOX2B^HA^ maintained its binding to PHOX2B/HLA-C*07:01 and PHOX2B/HLA-C*07:02 tetramers when expressed on primary T cells, and showed no binding to A*24 supertype alleles HLA-A*24:02 and HLA-A*23:01, in agreement with our structural findings and PepPred modeling (**Fig. 4b; Extended Data Fig. 21**). On-target pHLA tetramer staining by xTR-PHOX2B^HA^ is largely CD8 co-receptor independent, as indicated by similar levels of binding on CD8-positive versus CD8-negative Jurkat cells expressing the CAR (**Fig. 4c**). Upon co-culture with two HLA-C*07:01 PHOX2B-positive neuroblastoma cell lines SK-N-SH and Kelly, xTR-PHOX2B^HA^-CAR T cells demonstrated specific on-target killing at an effector-to-target (E:T) of 1:1, and no off-target killing using a HLA-C*07:02 positive, PHOX2B-negative colorectal cancer cell line SW620 (**Fig. 4d; Extended Data Fig. 22**). Notably, Kelly is an A9 serotype negative line, supporting the potential of xTR-PHOX2B^HA^-CAR for expansion of PHOX2B-targeted therapy to an orthogonal cohort of patients, relative to our existing 10LH PC-CAR^22,32^. We hypothesized that the reduced levels of killing were due to low antigen presentation, given the low HLA expression levels of neuroblastoma cells^33^, compounded by the reduced cell surface expression of HLA-C* alleles^34^. To enhance antigen density levels, we transduced neuroblastoma tumor cell lines with a single-chain trimer^35^ coupling the PHOX2B peptide to β_2_-microglobulin/HLA-C*07:02 heavy chain, and observed enhanced killing, supporting the notion that TRACeR-CAR T cell cytotoxicity is on-target, and can also cover HLA-C*07:02 positive cells (**Fig. 4e, Extended Data Fig. 23**). Furthermore, knockout of β_2_-microglobulin in the tumor cells abrogated killing, further supporting that the xTR-PHOX2B^HA^-CAR cytotoxicity observed for the parental cell lines is on-target and not due to bystander T cell activity **(Fig. 4e; Extended Data Fig. 23b)**.

**Figure 4.**
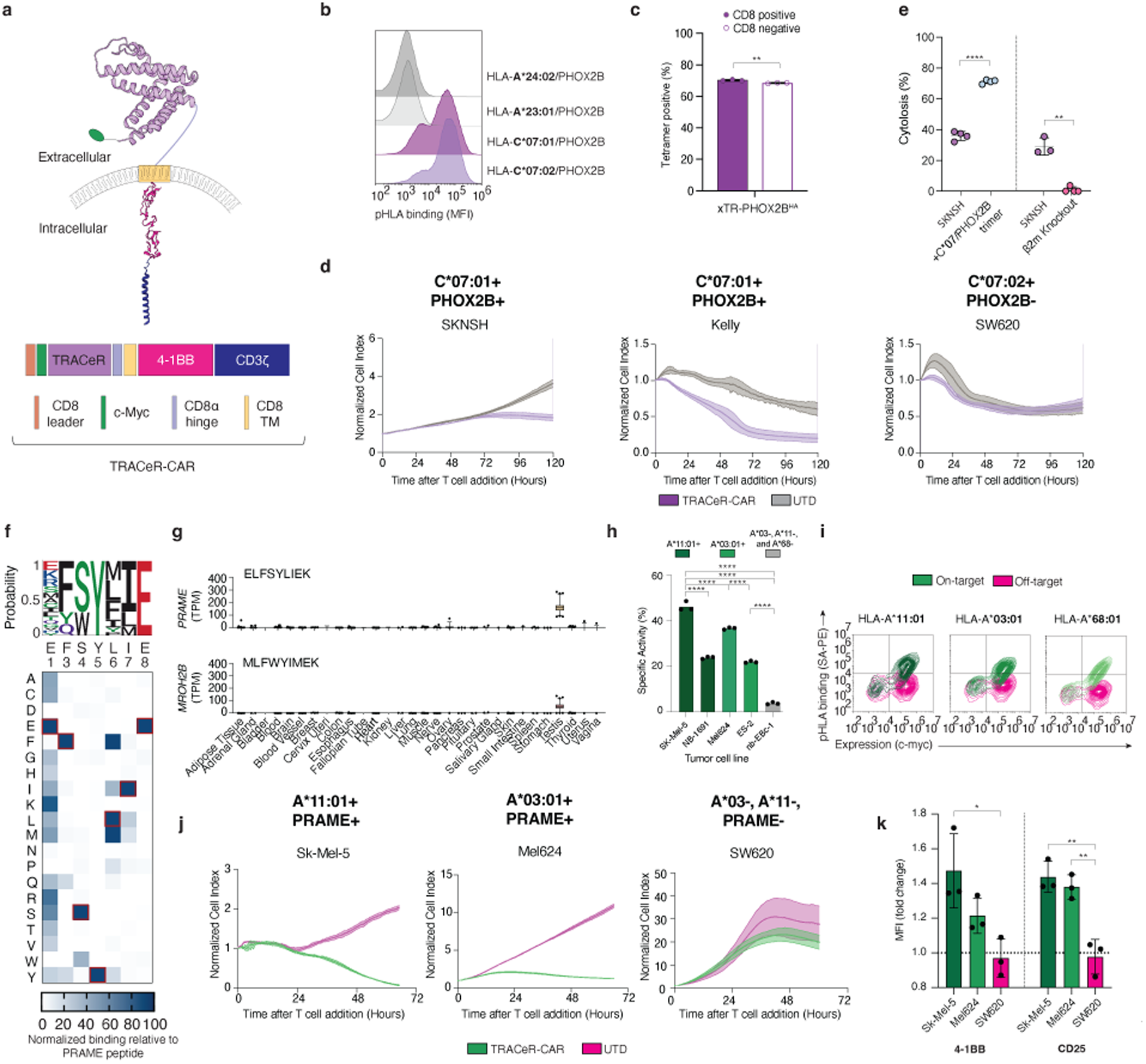
xTRACeR-CARs exhibit on-target cytotoxicity across HLA backgrounds *in vitro*. **a.** Schematic of xTRACeR incorporated into a second-generation CAR construct. **b.** pHLA tetramer staining of xTR-PHOX2B^HA^-CAR expressed on primary T cells. **c.** pHLA tetramer staining of xTR-PHOX2B^HA^-CAR expressed on CD8-positive versus CD8-negative Jurkat cells. **d.** Cytotoxicity of xTR-PHOX2B^HA^-CAR T cells co-cultured with two C*07:01 neuroblastoma cell lines and an off-target cell line at 1:1 E:T ratio. Data are shown as mean ± SD of 4 technical replicates and are representative of 2 independent experiments. **e.** Summary of cytotoxicity experiments performed as in (e) at 96 hours endpoint using SK-N-SH co-cultured with xTR-PHOX2B^HA^-CAR T cells, 1:1 E:T under either parental, enhanced PHOX2B pHLA expression levels using a single-chain PHOX2B- β_2_m-HLA-C*07:02 trimer, or β_2_m knockout. Percent cytolysis normalized to experiments using un-transduced (UTD) T cells. Data shown on the right and left come from independent experiments using different primary T cell donors **f.** X-scan of non-anchor PRAME peptide residues for binding to xTR-PRAME^HA^. Heatmap displays binding MFI relative to wild-type peptide, adjusted for background staining. Sequence logo displays amino acids with >15% relative binding MFI to WT (n=3). **g.** Tissue expression profiles of the *PRAME* and *MROH2B* genes, which contain the cognate (ELFSYLIEK) and potential off-target (MLFWYIMEK) peptide from our X-scan/ScanProsite^38^ search. We used The Genotype-Tissue Expression (GTEx) Portal^49^ to evaluate expression in healthy tissue, shown in TPM (Transcripts Per Million). **h.** Percent activation of NFAT-GFP reporter Jurkat cells expressing the xTR-PRAME^HA^-CAR, co-cultured with indicated tumor cell lines for 24 hours, 1:1 E:T. Specific activity = (GFP_Exp_ - GFP_Min_) / (GFP_Max_ – GFP_Min_); GFP_Min_ = unstimulated, GFP_Max_ = PMA-I. **i.** xTR-PRAME^HA^-CAR T cell co-staining by c-Myc and pHLA tetramers. **j.** Cytotoxicity of xTR-PRAME^HA^-CAR T cells co-cultured at 1:2 E:T ratio with PRAME-positive melanoma cell lines and a PRAME-negative off-target cell line, measured as in (e), n = 4. **k.** Fold change in MFI of T cell activation markers on CAR+ CD8+ T cells 24 h after co-culture in (k), normalized to xTR-PRAME^HA^-CAR T cells in media. Significance by unpaired t-test (c), unpaired t-test with Welch’s correction (e), and ordinary one-way ANOVA (h, k). ns, not significant, *p ≤ 0.05, **p ≤ 0.01, ***p ≤ 0.001, ****p ≤ 0.0001.

We next characterized the xTR-PRAME^HA^ peptide specificity landscape, and potential for cross-reactivity with off-target peptides from the self-ligandome. We used our chaperone-mediated peptide-exchange technology^36,37^ to generate a pHLA tetramer library containing all single amino acid substitutions of non-anchor PRAME residues on HLA-A*11:01. The tetramer library was used to probe binding to xTR-PRAME^HA^ displayed on yeast, revealing an overall highly target-specific profile that is tolerant for a limited scope of amino acid substitutions at positions 1, 3 to 4 and 6 to 7 (**Fig. 4f; Extended Data Fig. 24**). These results are in alignment with the peptide-centric antigen recognition mode and interactions observed in our cryo-EM structure (**Fig. 3b**). A ScanProsite^38^ search yielded one potential cross-reactive peptide (MLFWYIMEK) in the human proteome derived from *MROH2B*, which shows limited expression across healthy tissues (**Fig. 4g)**. However, a search across multiple published immunopeptidomics databases (IEDB^39^, HLA peptide ATLAS^40^, PCI-DB^41^) found no representation of the peptide in the ligandomes of healthy tissue. These results support a relevant therapeutic window for CAR-T development, which should be explored further during preclinical development.

We next used NFAT-inducible eGFP Jurkat (J^ASP90^) reporter cells^42^ to assess activation of xTR-PRAME^HA^ TRACeR-CARs and found specific activation upon co-culture with two HLA-A*11:01 and two HLA-A*03:01 expressing cancer cell lines, and minimal activation with an A03 supertype-negative cell line, demonstrating cross-allelic functional activity *in vitro* (**Fig. 4h and Extended Data Fig. 25**). The degree of activation correlates with HLA expression levels of each cell line, with the two melanoma cell lines (Mel624 and Sk-Mel-5) resulting in greater activation compared to the ovarian (ES-2) and neuroblastoma (NB-1691) lines (**Extended Data Fig. 26**). xTR-PRAME^HA^-CAR T cells maintained on-target binding to all three alleles and induced specific and potent tumor cell killing *in vitro* at an E:T of 1:2, accompanied by a corresponding increase in expression of activation markers 4-1BB and CD25 on CD8+ T cells (**Fig. 4i-k; Extended Data Fig. 27 and 28**). These results support that xTRACeRs can be deployed as stable antigen binding modules for CAR-T development.

Altogether, our work addresses a fundamental challenge shared by existing modalities targeting peptides derived from intracellular proteins, namely the HLA allotypic restriction which limits TCR- and antibody-based approaches to defined patient subpopulations. By combining computational prediction of conserved peptide presentation with cross-allelic binder engineering and CAR T cell validation, we show that this barrier can be overcome in a generalizable way. Structural analysis reveals that cross-allotype coverage and high peptide specificity are not mutually exclusive. Our engineered binders can achieve both by engaging conserved HLA residues through binding modes that can also navigate allotypic diversity. These results underscore a path toward off-the-shelf immunotherapies^43^ targeting public neoantigens and tumor-associated antigens across diverse patient populations^44,45^. Importantly, coverage of PRAME and PHOX2B antigens across 37–43% of the population suggests that a small library of xTRACeRs could enable supertype-stratified, off-the-shelf CAR-T therapies with near-universal reach.

The current study is limited by the number of allotypes examined, and future work will be needed to establish broader HLA compatibility. Our *in vitro* CAR-T results demonstrate efficacy across an HLA coverage pattern that complements existing PHOX2B PC-CAR^32^ and PRAME TCR-T^46^ therapeutics, supporting the potential for therapeutic expansion towards both targets. Our cross-reactivity analysis centered on the xTR-PRAME^HA^-CAR, guided by individual amino acid substitutions of the peptide, identifies a single potential off-target peptide, which is not represented in the self-ligandome. Nevertheless, further work will be needed to de-risk off-tumor toxicity arising from peptide cross-reactivity^47^. Validation in primary patient-derived tumor models *in vivo* remains an important next step, as does extending the platform to additional antigen targets of clinicial interest.

## Methods

### PepPred Development, Training, and Validation

Structures of 592 pHLAs from HLA3DB^12^ were first filtered by peptide antigen length. From 444 9mer peptide/HLA structures, we selected all cases where the same peptide is solved in complex with two different HLA alleles. This ground truth set of 104 pairs, comprised 86 individual pHLA structures covering 25 unique peptide antigens and 30 HLA allotypes. The peptide/HLA combinations in that set were run through a version of ^15^(AFF2) that has been finetuned ^48^ using all 9mer pHLA structures in HLA3DB^12^ to generate predicted models, and confidence metrics. Similarly, we used a finetuned version of Protpardelle^50^ which we trained on all pHLA structures. Each predicted pHLA structure was passed to condition Protpardelle all-atom sampling to generate 16 alternate peptide backbone structures for each pHLA target. Ensembles were generated using 80 rewind partial diffusion steps. Dihedral information from peptide residues 4-7 for each Protpardelle output decoy as well as the original AFF2 conditioning input were collected. To capture dominant variation, principal component analysis was performed independently for each given peptide of respective HLA allotypes, yielding 3D maps used to generate ensemble-level descriptors (see *Appendix*). Descriptive features were passed along with confidence scores and aggregate dihedral information and used to train an ensemble learner and produce predicted similarity scores using our ground truth set of 104 pHLA pairs. The model training and evaluation was done using 5-fold cross validation, leave one out cross validation, and label-shuffling to prevent data leakage and assess accuracy to unseen structural contexts.

### Identification of candidates for cross-allelic targeting

Initial sets of HLA alleles presenting established tumor-associated peptide antigens were obtained from the immune epitope database (IEDB)^39^. For each target antigen, the list of HLA-restricting alleles was expanded to include all other HLA alleles present within their respective HLA supertype^2^. The candidate HLAs were then evaluated for binding to the target utilizing NetMHCpan v4.2^17^, and all predicted strong or weak binders were advanced to the next step. Predicted binders with an allelic frequency greater than 0.05% in the US population from the NMDP database^51^ were retained for downstream analysis. Selected alleles were then evaluated using our AFF2/Protpardelle/PepPred pipeline to produce allele heatmaps clustered recursively to form a dendrogram.

### Yeast display and library screening

Yeast cells that were previously transformed were grown in SDCAA medium overnight at 30°C. Cells representing 10x coverage of the TRACeR library were then centrifuged at 2,400 *g* for 3 min and resuspended in SGCAA medium, supplemented with 0.2% glucose, at a cell density of 1×10^7^ cells per mL. Cells were then induced at 30°C for 20-24 h or at 20°C for 36-48 h. Cells to be used for staining were harvested and washed 1.5x times with PBSB (1x PBS and 1% BSA). Washed cells were labeled with biotinylated pHLA monomers or tetramers along with myc tag polyclonal chicken antibody (ThermoFisher A-21281). After incubation at 25°C for ∼1 h with shaking, cells were washed twice with ice-cold PBSB. Cells were then stained with goat-anti-chicken (H+L) secondary antibody Alexa fluorophore-488 (AF-488, ThermoFisher) for 10 min on ice. Following secondary staining, cells were washed twice with ice-cold PBSB and resuspended to be run on either the BD FACS Aria II cell sorter or the Sony SH800S cell sorter.

The theoretical diversity of our library was 1.7×10^10^ TRACeR variants. Thus, initial experimental screening using MACS was performed on 1×10^10^ yeast cells to maintain library coverage. For each target, one to two rounds of MACS and five to eight subsequent rounds of FACS were performed until diversity converged on clones matching our desired binding profiles. To confer cross-allelic profiles, toggling between orthogonal pHLA tetramers was initiated in round 3 of selections. NGS of final enriched pools was performed using AmpSeq Custom Sequencing services or Quintara Bio.

### Affinity maturation library preparation and cell selections

Plasmids of yeast clones isolated from naïve selections were extracted via mini-prep (Zymoprep Yeast Plasmid Miniprep II) and subjected to error-prone PCR using a mixture of 1x Taq Buffer (-MgCl_2_; ThermoFisher), 50mM MgCl_2_ (ThermoFisher), 10uM primers (Integrated DNA Technologies), 10mM dNTPS (ThermoFisher), 20uM 8-oxo-dGTP (ThermoFisher), 20uM dPTP ThermoFisher), 0.7ng template, and 0.05U/uL Taq DNA Polymerase (ThermoFisher) per manufacturers protocols. The yeast display plasmid, pCTCON2, was gapped using BamHI-HF and NheI-HF (New England Biolabs) and purified through gel extraction (QIAquick Gel Extraction Kit). Once sufficient insert and vector DNA were prepared, a starter culture of yeast strain EBY100 was grown overnight at 30°C in YPD medium. The following morning, cells were diluted in fresh, pre-warmed, YPD medium to a cell density of 5×10^6^ cells/mL and grown at 30°C with shaking. Once cells reached two-doublings (or a cell density between 1.5 and 2×10^7^ cell/mL), they were centrifuged at 2,400 *g* for 3 min. Cells were washed once more with sterile water and resuspended in 25 mL sterile water containing 100mM lithium acetate and 10mM DTT. Cells were incubated for 15 min at room temperature on a tube rotator and then washed twice with ice-cold sterile water. After the final wash, 1×10^9^ cells were aliquoted to Eppendorf tubes containing 1 ug of purified vector and 4 ug of purified insert and mixed thoroughly by vortexing. Cells were then transferred to pre-chilled 2 mm electroporation cuvettes. Using a BioRad gene-pulser XCell, competent yeast subjected to electroporation via a square wave protocol with the following parameters: 500 V, 15 ms pulse duration, 1 pulse, 2 mm cuvette size. Cells were recovered for one hour at 30°C before resuspension in SDCAA medium.

Once the library underwent two passages, cells representing 10x coverage of transformation efficiency were harvested and subjected to cell staining with pHLA tetramers or monomers. Generally, three to five rounds of FACS yielded TRACeR clones with at least a log-fold improvement in binding affinity over the parental naïve clone. Top-ranking clones matching our desired specificity profile were carried forward into downstream on-yeast and functional characterization.

### On-yeast titrations for affinity estimates

Individual yeast clones were grown in SDCAA medium overnight at 30°C. The following morning, cells were centrifuged at 2,400 *g* for 3 min and resuspended in SGCAA medium, supplemented with 0.2% glucose, at a cell density of 1×10^7^ cells per mL. Cells were then induced at 30°C for 20-24 h or at 20°C for 36-48 h. Cell to be used for staining were washed 1.5x times with PBSB and 2×10^6^ cells were aliquoted to corresponding wells in a 96-well microplate. Concurrently, monovalent streptavidin-PE (SA-PE) tetramers were generated via a 10-step serial addition of 0.1x SA-PE to a mixture containing a 3:1 molar ratio of soluble D-biotin to on- or off-target pHLA. Tetramerized antigens were serially diluted by 2x to produce a panel of an 11-point antigen titration for both on- and off-target pHLA reagents. Cells were then labeled with each concentration along with myc tag polyclonal chicken antibody for 1 h at 25°C with shaking. Following primary incubation, cells were washed 1.5x times with ice-cold PBSB and stained with goat-anti-chicken (H+L) secondary antibody Alexa fluorophore-488 (AF-488, ThermoFisher) for 10 min on ice. Upon completion of secondary staining, cells were washed twice with ice-cold PBSB and resuspended to be analyzed on the Cytoflex 3-laser flow cytometer or Bio-Rad ZE5 cell analyzer.

### Recombinant TRACeR expression and purification

DNA plasmids encoding the affinity matured TRACeR sequences were synthesized and cloned into pET-24a(+) *Escherichia coli* plasmid expression vectors (GenScript, N-terminal MBP and 6x-His tag). Plasmids were then chemically transformed into competent BL21 (DE3) *E. coli* cells. Single colonies were picked and a starter culture was grown in 2xYT medium, supplemented with 1% glucose, at 37°C overnight. The following morning, the cells were diluted 100-fold in fresh 2xYT medium, supplemented with 1% glucose, and grown at 37°C until the optical density (OD) reached 0.6 - 0.8. Protein expression was then induced with 1mM IPTG and grown overnight at 16°C. Induced cells were harvested and resuspended in 50 mM Tris buffer pH 8.0 and 150 mM NaCl, 0.5 mg/mL lysozyme, 0.1 mg/mL PMSF, and 0.1 mg/mL Benzonase. Lysis occurred for 1 h at room temp with slight agitation. The mixture was then sonicated and purified by nickel affinity followed by SEC (Superdex 75 10/300GL Increase, GE Healthcare).

### Recombinant peptide/HLA-I refolding, purification, validation, and tetramerization

DNA plasmids encoding the HLA-I heavy chains and human β_2_m-microglobulin (β_2_m) tagged with an N-terminal BirA substrate peptide (BSP, LHHILDAQKMVWNHR) were purchased from GenScript and transformed into competent BL21 *E. coli* cells (Novagen). BSP-tagged HLA-I I and β_2_m proteins were expressed in Luria-Bertani medium overnight and centrifuged for 20 minutes at 6,000 rpm. The cell pellet was resuspended in 12.5mL BugBuster media (Millipore), with the addition of 2mM phensylmethylosulfonyl fluoride, 10μL Benzonase Nuclease (Millipore), and 2.2uL rLysozyme (Millipore). The mixture was then incubated for 1 h at room temperature with slight agitation. Following incubation, cells were centrifuged at 10,0000 rpm for 15 minutes at 4°C. The inclusion body pellet was resuspended with 25mL TE Buffer (100mM Tris pH 8, 2mM EDTA) by sonication of 30 sec pulses at 40% amplitude and washed 2 subsequent times. Washed inclusion body pellets were resuspended in 6 mLs MES buffer (50mM MES, 0.1mM EDTA, pH 6.5). To the resuspended cell mixture, 4.8 g urea and 0.1mM fresh DTT were added before incubation for 1 h at room temperature with slight agitation. Solubilized inclusion bodies were then centrifuged at 6,000 rpm for 30 min at 4 °C. Refoldings were performed *in vitro* by diluting a 200 mg mixture of HLA-I and β_2_m at a 1:3 molar ratio in a step-wise manner in 1 L of refolding buffer (0.4M L-arginine 100 mM Tris pH 8, 2 mM EDTA, 4.9 mM reduced glutathione, and 0.57 mM oxidized glutathione) containing 10 mg of the desired peptide. pHLA refolding proceeded for 96 h at 4°C and subsequent protein purification via size-exclusion and ion-exchange chromatography was performed.

Following purification, pHLA protein stability was assessed via differential scanning fluorimetry (DSF). A fresh 50x dilution of SYPRO^TM^ Orange Protein Stain (5000x, ThermoFisher) was prepared in DSF Buffer (50mM NaCl, 20mM Na_3_PO_4_, pH 7.2). pHLA protein was added to a mixture containing 10x SYPRO^TM^ Orange Protein Stain to a final concentration of 10 μM in DSF buffer. The mixture was aliquoted in a MicroAmp Optical 384-well Reaction Plate (Biosystems), sealed with a MicroAmp Optical Adhesive film, and spun at 1000 rpm for 1 min. Once samples were readily prepared, they were subjected to a melt curve experiment starting at 25°C to 95°C in 0.017°C/s increments. The experiment was performed on a QuantStudio 5 real-time PCR machine with excitation and emission wavelengths set to 470 nm and 569 nm. Data analysis and fitting were performed in GraphPad Prism 9. Purified and quality-controlled protein was then biotinylated using the BirA500 biotin-ligase standard reaction kit (Avidity, LLC) per manufacturers protocols overnight at 4°C. Biotinylated product was washed 5 times the following day with 1x PBS pH 7.2 and flash frozen in the presence of 10% glycerol. Biotinylated protein was tetramerized via a 10-step addition of fluorophore-conjugated streptavidin to a mixture containing a 4:1 molar ratio of pHLA protein-to-streptavidin.

### Isothermal Titration Calorimetry

ITC experiments between PHOX2B/HLA-C*07:02 and TRACeR constructs were conducted using a MicroCal VP-ITC system (Malvern analytical). All proteins were exhaustively dialyzed into the buffer (150 mM NaCl, 20 mM sodium phosphate pH 7.2) and filtered through a 0.22 μm PES membrane. A syringe containing PHOX2B/HLA-C*07:02 at 150 μM with 0.45 mM peptide (QYNPIRTTF) was titrated into the calorimetry cell containing 5 μM TRACeR and the same peptide (0.45 mM). Injection volumes of 10 μL were performed for a duration of 10s and spaced 220s apart to allow a complete return to baseline. Data were processed and analyzed with Origin software. Isotherms were fit using a one-site ITC binding model. The first data point was excluded from the analysis. Reported K_D_, -T*ΔS, and ΔH values were determined using the 1-site binding model.

### Cryo-EM sample preparation

The TRACeR/pHLA complex was prepared by mixing xTR-PRAME^HA^ and PRAME (ELFSYLIEK)/HLA-A*11:01/β_2_m in 1:1 molar ratio and incubated on ice for 1 hour. Prior to the grid preparation, 0.04 % of Fluorinated Octyl Maltoside was added to the protein complex (3 mg/mL). Quantifoil R 1/2 300 Mesh, Cu grids were glow discharged at 15 mA for 60s. A sample volume of 3 µL was applied to the grids and subjected to vitrification using a Vitrobot Mark IV system (ThermoFisher) at 4°C with 100 % humidity. The sample was blotted for 3s after waiting time of 5s and a blot force of 2 using Standard Vitrobot 595 filter paper (Ted Pella, Inc). The grids were plunge-frozen into liquid-nitrogen-cooled liquid ethane. The same protocol was followed to prepare grids for xTR-PHOX2B^HA^/PHOX2B/HLA-C*07:02/ β_2_m complex, except 0.08 % of Fluorinated Octyl Maltoside was added to the protein complex (4 mg/mL).

### Cryo-EM data collection, processing, and model building

Cryo grids of the complexes were imaged at 130,000× nominal magnification using a Falcon 4 detector (ThermoFisher) on a Glacios 2 (ThermoFisher) microscope operating at 200 kV with a calculated pixel size of 0.887 Å. Automated image collection was performed using EPU with a nominal defocus range of –0.8 to –2.1 µm. For xTR-PRAME^HA^ /PRAME/HLA-A*11:01/β_2_m complex, 5,325 movies were collected. Data processing for all datasets was carried out in CryoSPARC v4.7.1. Micrographs were aligned using Patch-motion correction and the Contrast Transfer Function (CTF)-corrected using Patch CTF Estimation in CryoSPARC. Junk micrographs were removed using Manually Curate Exposures tool. Blob picker was used to pick particles from a subset of 1,234 micrographs with a minimum particle diameter of 80 Å to a maximum of 140 Å. An initial round of 2D classification was used to generate templates for template picker. A total of 1,001,606 particles were extracted from all micrographs after template picker. A round of 2D classification were performed to remove junk particles. This yielded 323,234 particles which were then subjected to ab-initio reconstruction to generate five volumes. We also selected best 2D classes to generate a volume as a “good” reference for heterogeneous refinement. Four rounds of heterogeneous refinements were performed using particles extracted from all micrographs (1,001,606 particles) and three volumes as references (one “good” and two “bad” volumes from ab-initio reconstructions). The resulting volume and 130,398 particles corresponding to it were used for non-uniform refinement which yielded a 3.96 Å resolution map. For xTR-PHOX2B^HA^/PHOX2B/HLA-C*07:02/β_2_m complex, 5,842 movies were collected. Micrographs were aligned using Patch-motion correction and the Contrast Transfer Function (CTF)-corrected using Patch CTF Estimation in CryoSPARC. Micrographs with CTF fit worse than 10 Å were discarded. Blob picker was then used to pick particles from entire dataset with minimum particle diameter of 80 Å to a maximum of 140 Å. A round of 2D classification was performed to remove junk particles. This yielded 493,760 particles which were then subjected to ab-initio reconstruction to generate three volumes. Meanwhile, best looking 2D classes were used as templates to pick 4,288,572 particles from the entire dataset. Four rounds of heterogeneous refinements were performed using all particles (4,288,572 particles) and the three ab initio volumes. The final best set of 517,824 particles was used for non-uniform refinement which yielded a 3.89 Å resolution map. For model building, initial models for individual TRACeRs and peptide/HLAs were generated using AlphaFold 3 and fitted into the respective density maps using UCSF Chimera. The models were then manually adjusted in Coot followed by real-space refinements in PHENIX package.

### Peptide X-scan assays

For tetramer library preparation, streptavidin–(R)-phycoerythrin (PE) (Agilent Technologies) was added to open-HIV/HLA-A*11:01 at a 4:1 molar ratio of pHLA monomer to streptavidin protein every 10 min over ten intervals at room temperature in the dark. Concurrently, peptide concentrations were calculated using the Pierce Quantitative Fluorometric Peptide Assay (ThermoFisher) and analyzed in GraphPad Prism 10. Tetramerized open-HIV/HLA-A*11:01 was then mixed with TAPBPR (7:1 molar ratio of open-HIV-loaded HLA-A*11:01 to TAPBPR) and individual peptides from a site-saturated mutagenesis library of PRAME peptide positions 1, 3, 4, 5, 6, 7, and 8 (1:10 molar ratio of tetramerized HIV-loaded HLA-A*11:01 to peptide; ElimBio). Each reaction was incubated 3 h at 32°C followed by overnight at 4°C. Tetramerized and peptide-exchanged proteins were washed using Amicon Ultra centrifugal filter units with a 100-kDa membrane cutoff to remove excess peptides and TAPBPR with a 1:1,000 dilution of 1x PBS buffer. The resulting tetramers could be stored at 4 °C for up to 4 weeks.

Yeast cells encoding xTR-PRAME^HA^ were grown in SDCAA medium overnight at 30°C and induced in SGCAA medium for 20-24 h at 30°C. Induced cells were washed 1.5x times with ice-cold PBSB and 2×10^6^ cells were aliquoted to corresponding wells in a 96-well microplate. Cells were then labeled with each peptide-exchanged tetramer from the SSM library along with myc-tag polyclonal chicken antibody (ThermoFisher) for 30 min at 25°C with shaking. Following primary incubation, cells were washed 1.5x times with ice-cold PBSB and stained with goat-anti-chicken (H+L) secondary antibody Alexa fluorophore-488 (AF-488, ThermoFisher) for 10 min on ice. Upon completion of secondary staining, cells were washed twice with ice-cold PBSA and resuspended to be analyzed on the Cytoflex 3-laser flow cytometer.

Individual peptides from a site-saturated mutagenesis library of PHOX2B peptide positions 1, 3, 4, 5, 6, 7, and 8 were reconstituted per manufacturer protocols (Mimotopes). Peptide concentrations were calculated using the Pierce Quantitative Fluorometric Peptide Assay (ThermoFisher) and analyzed in GraphPad Prism 10. Each peptide was pulsed on C*07:02 monoallelic B721.221 cells (at 10uM) for 2 h at 37°C. Peptide pulsed cells were then stained with 1ug/mL of soluble tetramerized xTR-PHOX2B^HA^ conjugated with streptavidin–(R)-phycoerythrin (PE) (Agilent Technologies) at a ratio of 4:1. Stained cells were washed twice with ice-cold FACS buffer (1x DPBS, 2.5% heat-inactivated fetal bovine serum FBS, 0.02% sodium azide) and resuspended to be analyzed on the Cytoflex 3-laser flow cytometer.

### Lentiviral production

One day prior to transfection, Lenti-X 293T cells (Takara Bio) were cultured in DMEM (Gibco), 10% heat-inactivated FBS (Seradigm), and 1% Glutamax (Gibco). When cells reached a confluency between 70-80%, cells were transfected with Lipofectamine (Thermo Fisher Scientific) using pMD2.G (Addgene #12259), psPAX2 (Addgene #12260), and transfer plasmid (pTRPE vector from Carl June lab for CAR constructs). The media containing virus was harvested at 24, 48, and 72-hours post-transfection. The media was clarified by centrifuging at 500xg for 10 minutes before adding 1/3 volume of Lenti-X Concentrator (Takara Bio). All harvests were pooled and incubated for 1 hour to overnight before concentration of virus. Virus containing media was then concentrated by centrifugation at 1500xg, 4 °C for 45 minutes and the resulting pellet concentrated 50-100X by resuspension in 1X cold PBS. Virus was then either used immediately or aliquoted, flash frozen in liquid nitrogen and stored at −80 °C.

### General cell culture

B721.221 monoallelic cells were generously provided by Dr. Derin Keskin at Harvard University. J^ASP90^ CD8^+^/TCRαβ null Jurkat E6.1 cells engineered to express NFAT-inducible eGFP were generously provided by Dr. Beatriz M. Carreno and Dr. Gerald P. Linette from the University of Pennsylvania. All tumor cell lines were provided by Dr. John M. Maris at the Children’s Hospital of Philadelphia. The following cell lines were cultured in R10 media, comprising of RPMI 1640 (Gibco), 10% heat-inactivated FBS (Seradigm), 1% 100x GlutaMAX (Gibco), 1% 10,000 U/mL penicillin/streptomycin (Gibco), and 1% 1M HEPES (Gibco): B721.221 cells, Jurkat cells, human T cells, SK-N-SH, SW620, Mel624, NB1691, nb-EBc-1. ES-2 cells were cultured in DMEM (Gibco), 10% heat-inactivated FBS (Seradigm), 1% 100x GlutaMAX (Gibco), 1% 10,000 U/mL penicillin/streptomycin (Gibco), and 1% 1M HEPES (Gibco). Sk-Mel-5 cells were cultured in EMEM (Gibco), 10% heat-inactivated FBS (Seradigm), 1% 100x GlutaMAX (Gibco), 1% 10,000 U/mL penicillin/streptomycin (Gibco), and 1% 1M HEPES (Gibco). All cells are maintained at 37°C, 5% CO_2_. SK-N-SH cells were further transduced with single-chain trimer consisting of PHOX2B peptide coupled to β_2_-microglobulin and HLA-C*07:02 heavy chain with mKate2 NLS expression.

### CRISPR-Cas9 editing

Neuroblastoma cells were edited by electroporation of CRISPR-Cas9 ribonucleoproteins (RNPs) to knock out β_2_-microglobulin. Cas9 was complexed with β_2_m-specific single guide RNAs (sgRNA) 5’-CGGAGCGAGAGAGCACAGCG-3’, 5’-GGCCGAGAUGUCUCGCUCCG-3’, and 5’-ACUCACGCUGGAUAGCCUCC-3’ (Synthego) at a 1:3 ratio of Cas9/sgRNA and electroporated using a Lonza 4D-Nucleofector X Unit program EH-100 and buffers from P3 primary cell 4D-nucleofector X Kit (Lonza). RNPs were prepared by adding in the following order: 5 μl of P3 buffer, 1.2 μl of sgRNA (100 μM), and 2 μl of Cas9 (20 μM) per reaction. An amount of 8.2 μl of this mixture was added to 200,000 cells resuspended in 15 μl of P3 buffer. Reactions were performed in 16-well strips.

### Jurkat production and activation assays

Jurkat cells were transduced with TRACeR-CAR lentivirus and sorted to purity based on c-Myc expression. To test for antigen specific activation, sorted TRACeR-CAR Jurkat cells were mixed at 1:1 with 40,000 tumor cells plated on the previous day. After 24 h, cells were analyzed by flow cytometry to determine the percentage of GFP-expressing cells in each sample. Cells activated with PMA (50 ng/mL) and ionomycin (750 ng/mL) were incubated as positive controls.

### CAR T cell production

Freshly isolated CD4 and CD8 T cells were mixed in a 1:1 ratio, cultured in R10 media. T cells were cultured at 1 million cells/mL and activated with a 3:1 ratio of Dynabeads Human T-Activator CD3/CD28 beads (Gibco) for 24 hours before addition of lentivirus. T cells were stimulated with anti-CD3/CD28 beads for a total of 72 hours before de-beading and maintained at a density of 0.8 x 10^6^ cells per mL for 12 days. Lentiviral transduction efficiency was determined by co-staining with PE-conjugated pHLA tetramers and anti-FITC c-Myc antibody (Miltenyi Biotec, 130-116-485).

### In vitro CAR T efficacy studies

CAR T cell cytotoxicity was monitored in real time using an impedance-based assay (xCELLigence RTCA system, Agilent). Cells were seeded per well in 96-well E-plates, and CAR T cells were added to the wells at different effector-to-target (E:T) ratios after 24 h. Electrode impedance was recorded every 15 minutes and expressed as a dimensionless Cell Index (CI). For data analysis, cell index was normalized to the time point of CAR T cell addition.

### CAR T flow cytometry

For flow cytometry, cells were stained in fluorescence-activated cell sorting (FACS) buffer consisting of 1x DPBS, 2.5% heat-inactivated fetal bovine serum, and 0.02% sodium azide. The following antibodies along with vendor, catalog number, and working dilution were used: Near IR (876) fluorescent reactive dye (Thermo Fisher, L34982A, 1:1000), anti-human CD8-FITC (BioLegend, 980908, 1:100), anti-human CD4-APC (BioLegend, 300514, 1:100), anti-human LAG-3-PerCP-eFlour 710 (Thermo Fisher, 46-2239-42), anti-human CD279 PD-1-BV421 (BioLegend, 329920, 1:100), anti-human CD366 Tim-3-PE/Cyanine7 (BioLegend, 345014, 1:100), anti-human CD25-PE/Dazzle 594 (BioLegend, 385224, 1:100), anti-human CD137 4- 1BB-PE/Cyanine7 (Biolegend, 300808, 1:100), anti-human CD69-BV421 (BioLegend, 310930, 1:100), anti-human c-Myc-FITC (Miltenyi Biotec, 130-116-653, 1:50), anti-human c-Myc-PE (Miltenyi Biotec, 130-126-812, 1:50). Anti-human c-Myc-Alexa Fluor 647 (BioLegend, 626810, 1:50). Samples were acquired on a Cytoflex 6L or 3L, and data analyzed with FlowJo v10 software (FlowJo, LLC).

### Ethical approval and informed consent

The Human Immunology Core at the University of Pennsylvania provided all primary cells used in this study. The studies involving human participants were reviewed and approved by the University of Pennsylvania Institutional Review Board (IRB# 850312). A written informed consent to participate in this study was provided by the participants.

### Data, Code, and Materials Availability

All data needed to evaluate the conclusions in the paper are present in the paper and/or the Extended Data Materials. Plasmid DNA encoding the TRACeR construct developed in this work can be obtained by the authors upon reasonable request. A fully functional code for running the PepPred pipeline can be obtained from https://zenodo.org/records/20315909.

Finetuned model parameters for PepPred, AFF2, and Protpardelle can be obtained from https://zenodo.org/records/20076767. Atomic coordinates and cryoEM densities for the PRAME and PHOX2B complexes have been deposited in the PDB under the accession codes 13FG and 13FH.

## Acknowledgements

The authors thank Dr. Beatriz M. Carreno and Dr. Gerald P. Linette (University of Pennsylvania) for providing the J^ASP90^ reporter cells, Emily Cento, Zhilin Chen, Max A. Eldabbas, and Emileigh Maddox of the Human Immunology Core and the Division of Transfusion Medicine and Therapeutic Pathology at the Perelman School of Medicine at the University of Pennsylvania (RRID: SCR_022380) for providing de-identified primary T cells, and the Institute of Structural Biology and Beckman Center for cryo-EM at the University of Pennsylvania (RRID: SCR_022375) for assistance with cryo-EM data collection.

## Funding

This work was delivered as part of the NexTGen Team supported by the Cancer Grand Challenges partnership funded by Cancer Research UK (CGCATF-2021/100002), the National Cancer Institute (CA278687-01), and The Mark Foundation for Cancer Research. We acknowledge additional support by NIH grants from NIGMS (7R35GM125034 to N.G.S.; R01GM147893 to P.-S.H.) and NCI (R35CA220500 to J.M.M.), a Cancer Early Detection award by The Mark Foundation for Cancer Research (to N.G.S, P.-S.H) and American Cancer Society grant (ACS134055-IRG-218 to P.-S.H). E.S. was supported through a sponsored research agreement with Notamab Therapeutics Inc., and an award from the Lindonlight Collective. D.H. was supported by an NCI T32 training grant (5-T32-CA-009140-50). The Human Immunology Core at the Perelman School of Medicine at the University of Pennsylvania is supported in part by NIH (P30 AI045008 and P30 CA016520).

## Author contributions

Writing of the original draft was performed by W.B., E.L.S., S.M.S., O.S. P.-S.H., and N.G.S. Conceptualization of the experiments was performed by W.B., E.L.S., S.M.S., O.S., C.H.J., R.M.Y., J.M.M., P.-S.H., and N.G.S. Investigation was performed by W.B., E.L.S., S.M.S., O.S., R.P., S.J.W., P.R., H.D., Y.S., D.H., C.S.W., T.L., E.L., R.K., M.Y.W., P.-S.H., and N.G.S. The review and editing of the manuscript was performed by W.B., E.L.S., S.M.S., O.S., C.H.J., R.M.Y., J.M.M., P.-S.H., and N.G.S. Methodology was formulated by W.B., E.L.S., S.M.S., O.S., R.P., S.J.W., P.R., H.D., Y.S., D.H., C.S.W., T.L., E.L., R.K., M.Y.W., P.-S.H., and N.G.S. Resources were provided by C.H.J., R.M.Y., J.M.M., P.-S.H., and N.G.S. Data curation was performed by W.B., E.L.S., S.M.S., O.S. Validation was performed by W.B., E.L.S., S.M.S., and O.S. The project was supervised by C.H.J., R.M.Y., J.M.M., P.-S.H., and N.G.S. Project administration was carried out by P.-S.H., and N.G.S. Visualizations were developed by W.B., E.L.S., S.M.S., and O.S. Funding acquisition was performed by C.H.J., R.M.Y., J.M.M., P.-S.H., and N.G.S.

## Competing interests

W.B., E.L.S., P-S.H. and N.G.S. are inventors on a patent application related to this work. P-S.H. and N.G.S. are scientific co-founders and advisors to Notamab Therapeutics. The Sgourakis laboratory receives research funding from Notamab Therapeutics in the form of a sponsored research agreement. The remaining authors declare no competing interests.

