## Extended Data Figures, Tables, and Appendix for "A generalizable system for antigenic peptide targeting across HLA-I allotypes"

### Extended Data Figures and Tables

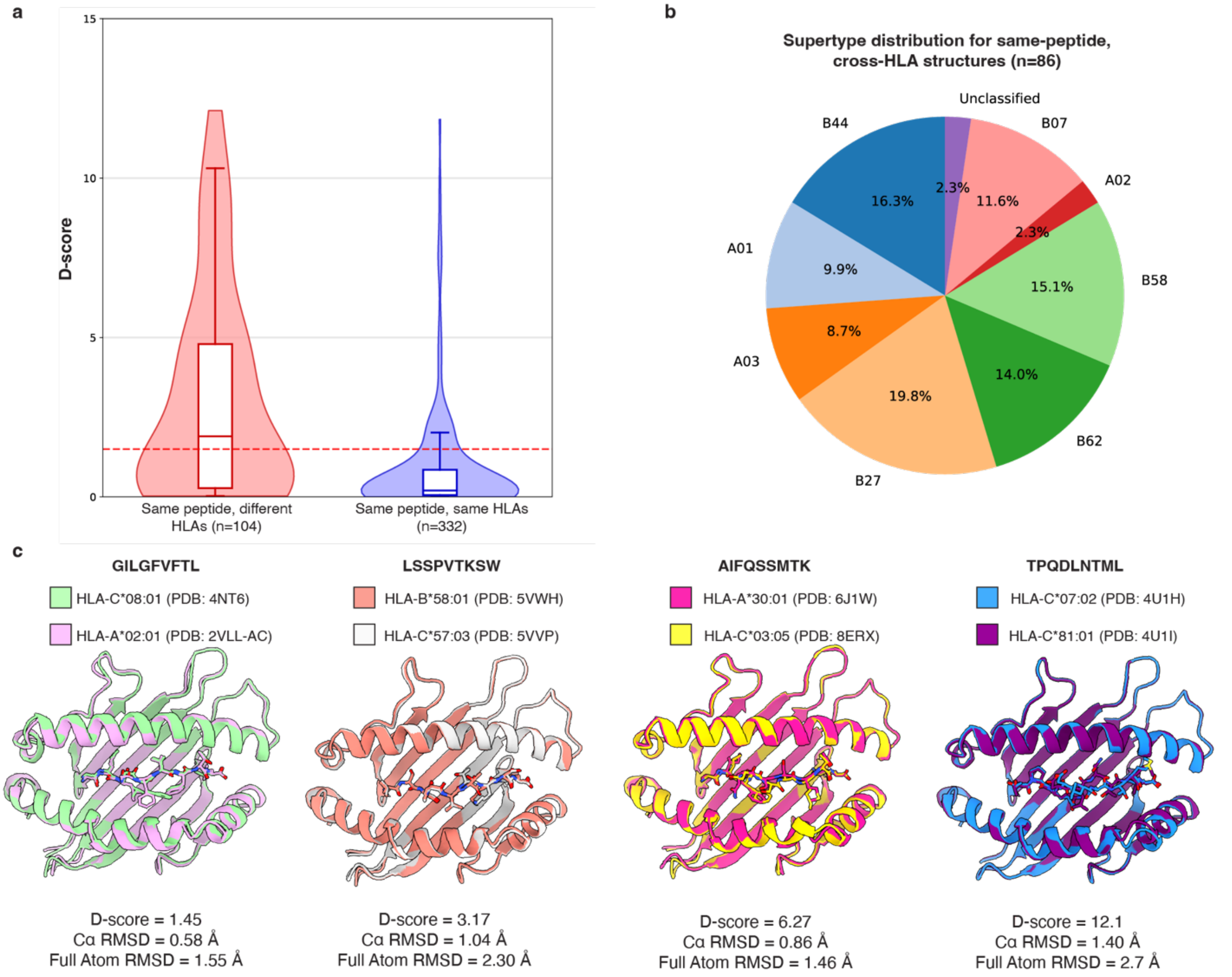


**Extended Data Fig. 1. Analysis of peptide conformational divergence across HLA allotype pairs of known structures. a.** D-scores for select pairs of 9mer peptide/HLA structures. For cases where the same peptide is presented across two different HLAs (termed cross-HLA), 46% of structures exhibit a D-score below 1.5, which are defined as conformationally similar. When the same peptide/HLA complex has been captured in independent X-ray structures or in independent copies within the asymmetric unit of a single X-ray structure, we observe consistent conformations with nearly all D-scores below 1.5. **b.** Distinct cross-HLA structures are distributed across HLA-A* and -B* supertypes. **c.** Select cross-HLA structural examples. D-score readily increases as peptide conformations diverge, more effectively capturing conformational variation than with traditional RMSD metrics.

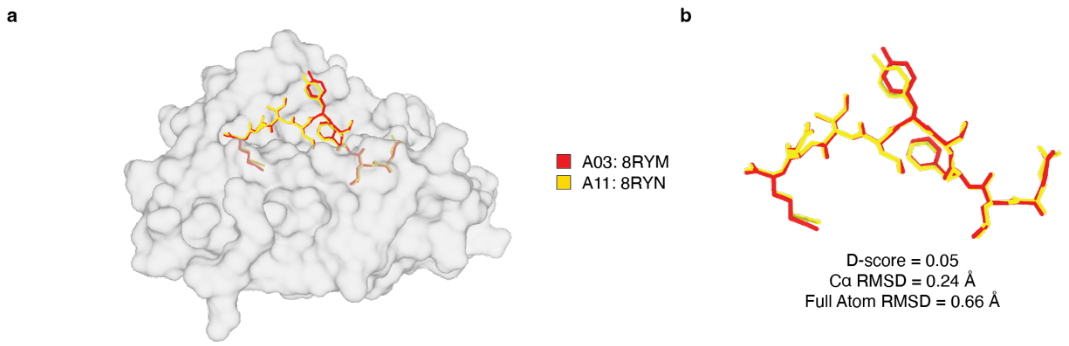


**Extended Data Fig. 2. The PRAME ELF peptide is presented in a conserved conformation on HLA-A*03:01 and HLA-A*11:01 according to solved structures a.** Overlay of X-ray structures containing a PRAME epitopic peptide (ELFSYLIEK) presented by HLA-A*03:01 (PDB: 8RYM) and HLA-A*11:01 (PDB: 8RYN). **b**. Extracted peptide conformations of PRAME bound to HLA-A*03:01 (red) and HLA-A*11:01 (yellow) with a D-score of 0.05 / RMSD of 0.66 Å, indicating high structural similarity.

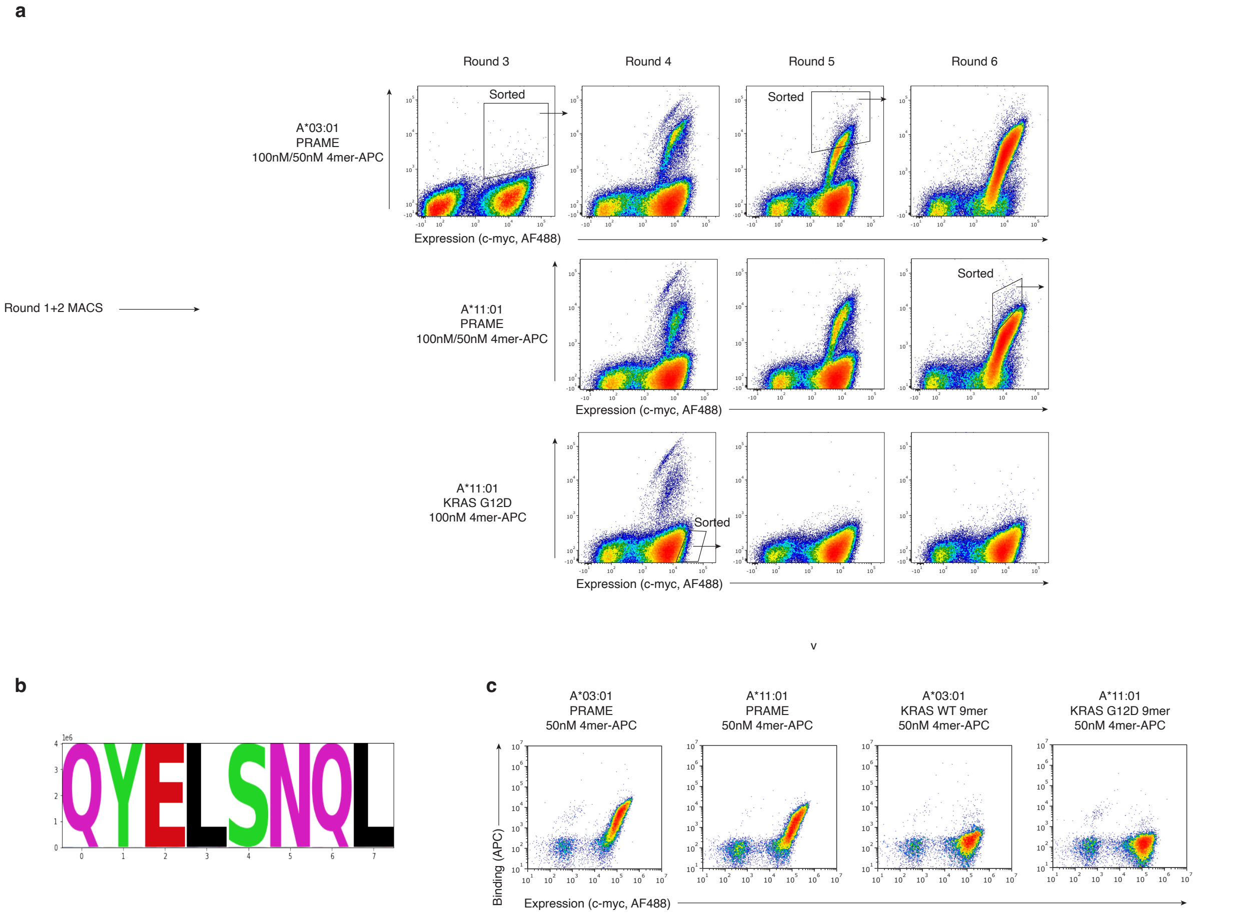


**Extended Data Fig. 3. Development of xTR-PRAME. a.** Enrichment process starting from naïve TRACeR library using tetramerized pHLA targets containing the ELFSYLIEK peptide bound to HLA-A*03:01 and HLA-A*11:01. **b.** Logo plots of the TRACeR specificity box (diversified in the naïve TRACeR library) from enriched libraries after sorting against both targets**. c.** On-yeast pHLA tetramer staining of enriched xTR-PRAME clone showing binding to the target peptide, and no binding to irrelevant peptides presented on both HLA allotypes.

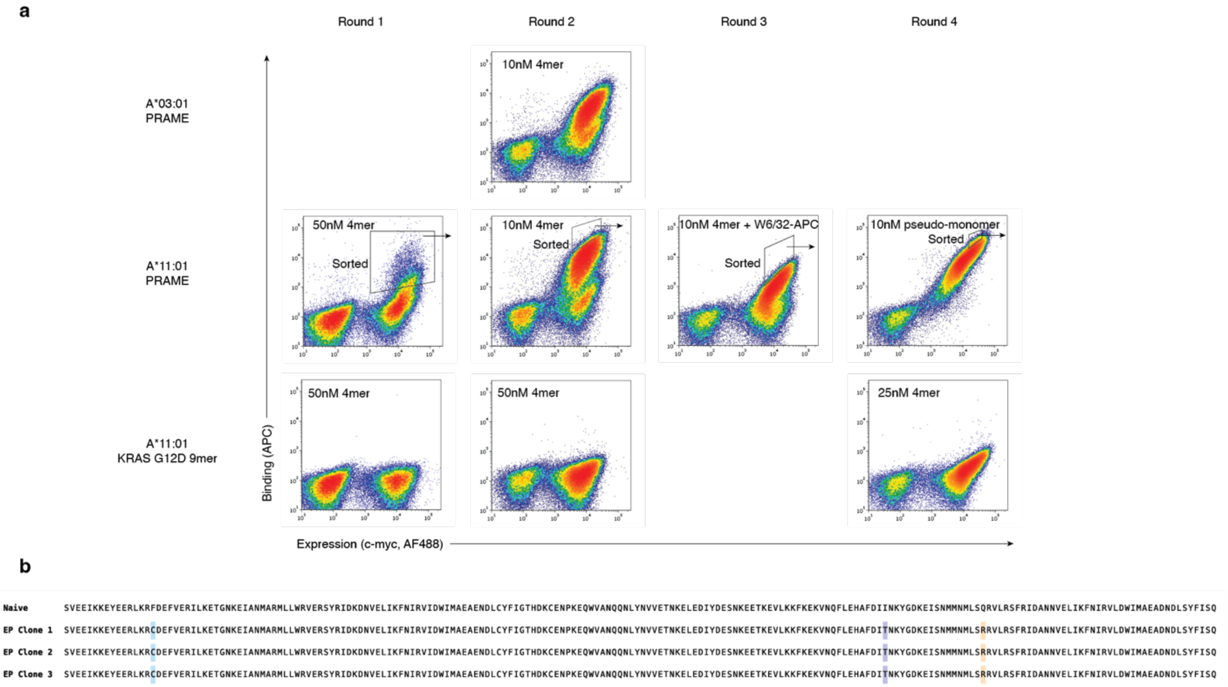


**Extended Data Fig. 4. Development of xTR-PRAME$^{HA}$ using error-prone mutagenesis. a.** Enrichment process for affinity-matured xTR-PRAME. **b.** Sequence alignment of colonies from R4 sort demonstrate sequence convergence.

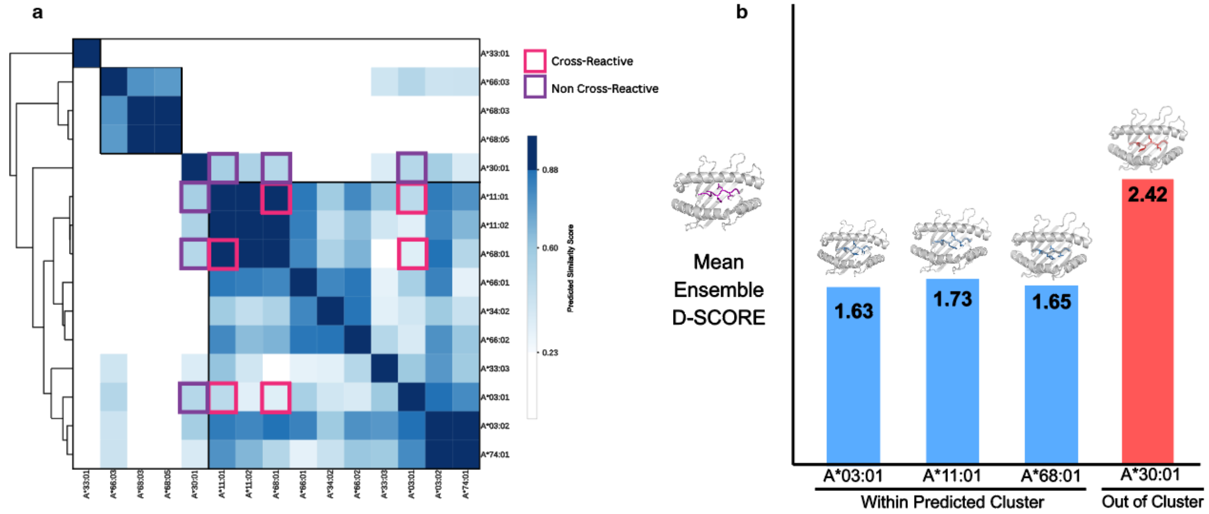


**Extended Data Fig. 5. Conformational analysis of PRAME peptide presentation across HLAs. a.** PepPred structural similarity scores are plotted for all HLA pairs from the A*03-A*11 supertype. Alleles confirmed as Strong or Weak Binders via NetMHCpan were compared for structural similarity using PepPred with inputs from Protpardelle and AlphaFold-FineTune. Cells comparing confirmed cross-reactive pHLA pairs are outlined in magenta, whereas confirmed non-reactive pairs are outlined in purple. **b.** Mean D-Score between the predicted structural ensembles for PRAME by Protpardelle & AlphaFold FineTune and solved crystal structure of PRAME presented on A*11:01 (PDB: 8RYN). Overlayed pHLA structures of the predicted ensembles for are shown for A*03:01, A*11:01, A*68:01, and A*30:01 and colored and annotated based on PepPred predicted structural relationship to the solved pHLA structure as Predicted Cluster (Blue) or Out of Cluster (Red). The pHLA dimer of the solved crystal structure of PRAME presented on A*11:01 (PDB: 8RYN) is shown in purple.

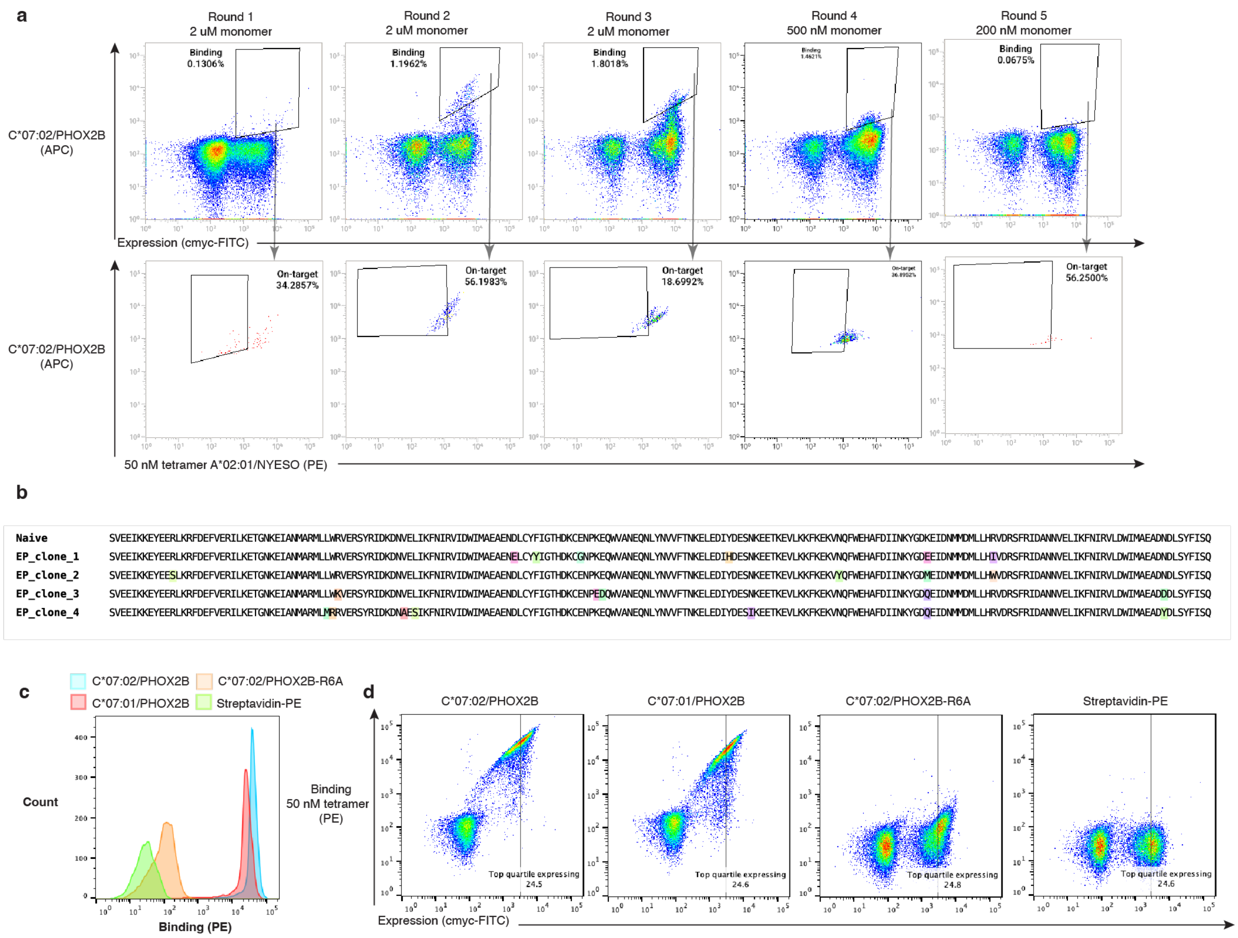


**Extended Data Fig. 6. Development of xTR-PHOX2B$^{HA}$ using error-prone mutagenesis. a.** Enrichment of high-affinity clones binding to C*07:02/PHOX2B (QYNPIRTTF) by iterative rounds of cell sorting with decreasing concentrations of monovalent pHLA reagent, starting from our previously reported naïve binder. 50 nM PE tetramer of A*02:01/NYESO (SLLMWITQV) was used as the off-target negative control. Each round, the expressing population is gated for binding signal and further gated against APC signal from staining with the off-target reagent. Cytometry plots show sorts from a sample of 100,000 cells and are representative of a longer total sort. **b.** Colony picking yielded four clones, which show multiple mutations of K149. EP_clone_4 was carried forward as xTR-PHOX2B$^{HA}$. **c.** Histogram showing xTR-PHOX2B$^{HA}$ binding to on-targets C*07:02/C*07:01/PHOX2B but not C*07:02/PHOX2B-R6A peptide (QYNPI**A**TTF). **d.** Cytometry plots of xTR-PHOX2B$^{HA}$ binding to targets in c.

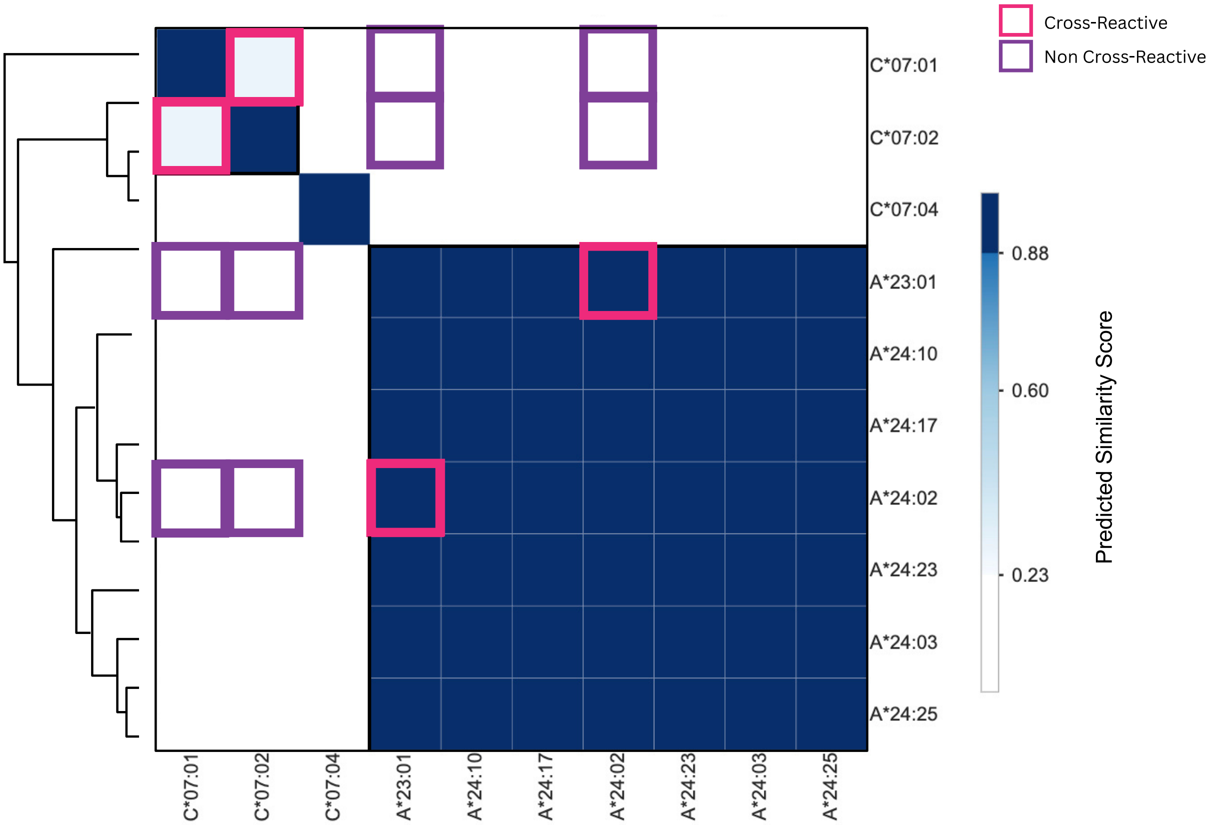


**Extended Data Fig. 7. Conformational analysis of PHOX2B peptide presentation across HLAs.** PepPred structural similarity scores are plotted for all HLA pairs from the A*24 and C*07 supertypes. Alleles with confirmed binding to PHOX2B were compared for structural similarity using PepPred with inputs from Protpardelle and AlphaFold-FineTune. Pairs of HLA alleles with confirmed cross-reactivity for binders in the present study (C*07:01/C*07:02) or our previous study (A*24:02/A*23:01) are outlined in magenta. Purple squares indicate HLA allelic pairs with no possible cross-reactivity for binders due to divergence in the conformation of the PHOX2B peptide.

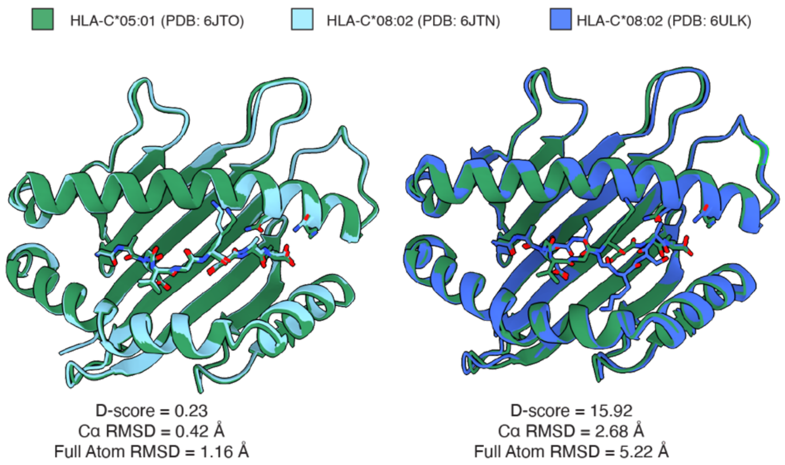


**Extended Data Fig. 8. Structural conservation of the G12D peptide on HLA-C*05:01 and HLA-C*08:02.** Analysis of X-ray structure containing the KRAS-G12D peptide (GADGVGKSAL) bound to HLA-C*05:01 (PDB: 6JTO) and HLA-C*08:02 (PDB: 6JTN, 6ULK). Polymorphic residues between C*08:02 and C*05:01 near the peptide C-terminus are shown as sticks. Two independent structures of KRAS-G12D peptide presented on HLA-C*08:02 observe distinct peptide conformations, one of which overlaps with HLA-C*05:01 structure (D-score = 0.23 for 6JTO/6JTN vs 15.92 for 6JTO/6ULK).

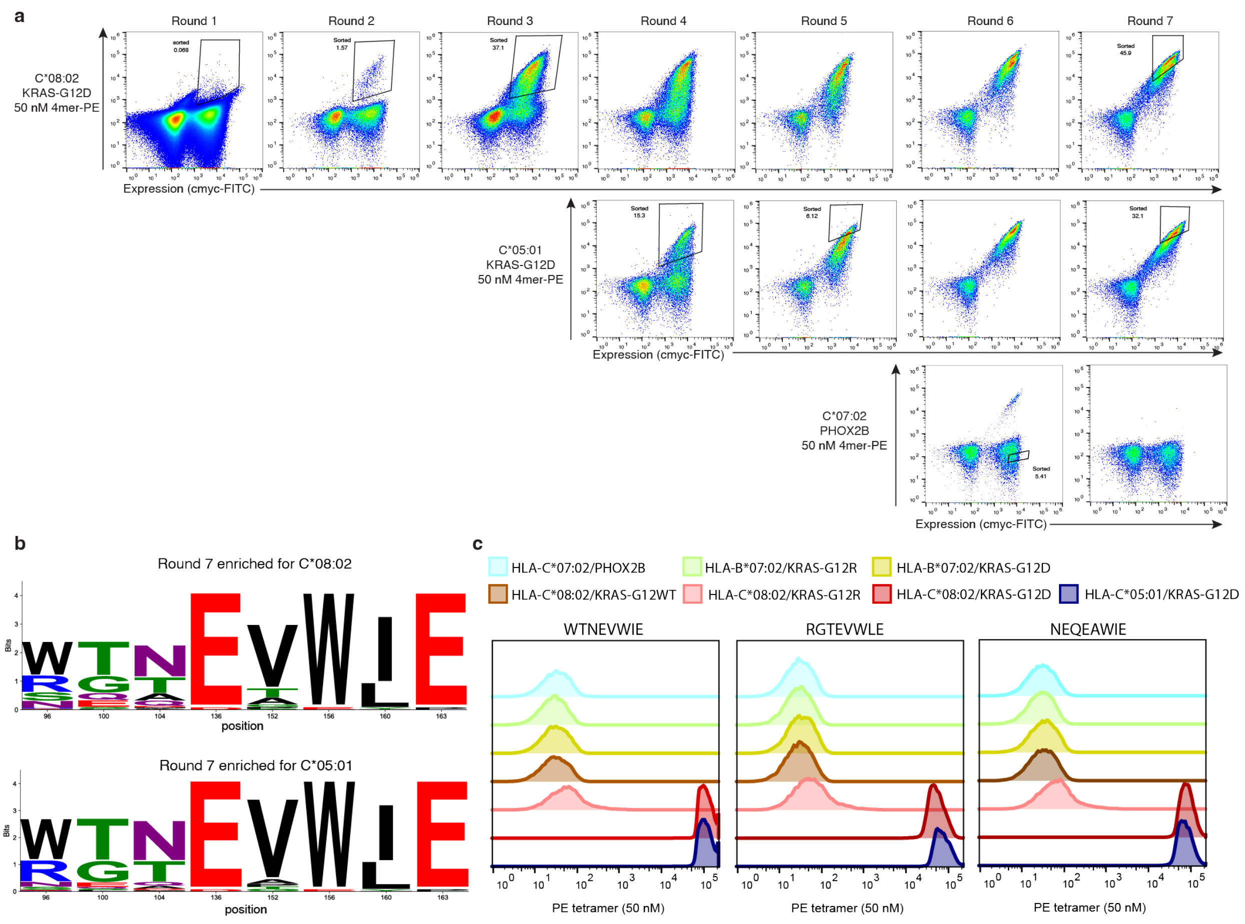


**Extended Data Fig. 9. Development of xTR-KRAS. a.** Naïve enrichment of TRACeR library against HLA-C*08:02 and HLA-C*05:01 presenting KRAS-G12D (GADGVGKSAL). **b.** Logo plots of the TRACeR specificity box from enriched libraries after sorting against both HLA-C*08:02/KRAS-G12D and HLA-C*05:01/KRAS-G12D. **c.** On-yeast specificity of top three clones ranked by sequencing counts. All three TRACeRs show specific binding to both HLA-C*08:02 and HLA-C*05:01 presenting the KRAS-G12D peptide but not the KRAS-WT peptide nor other KRAS antigens presented by off-target alleles. The top clone containing WTNEVWIE sequence was carried forward through affinity maturation, hereafter referred to as xTR-KRAS.

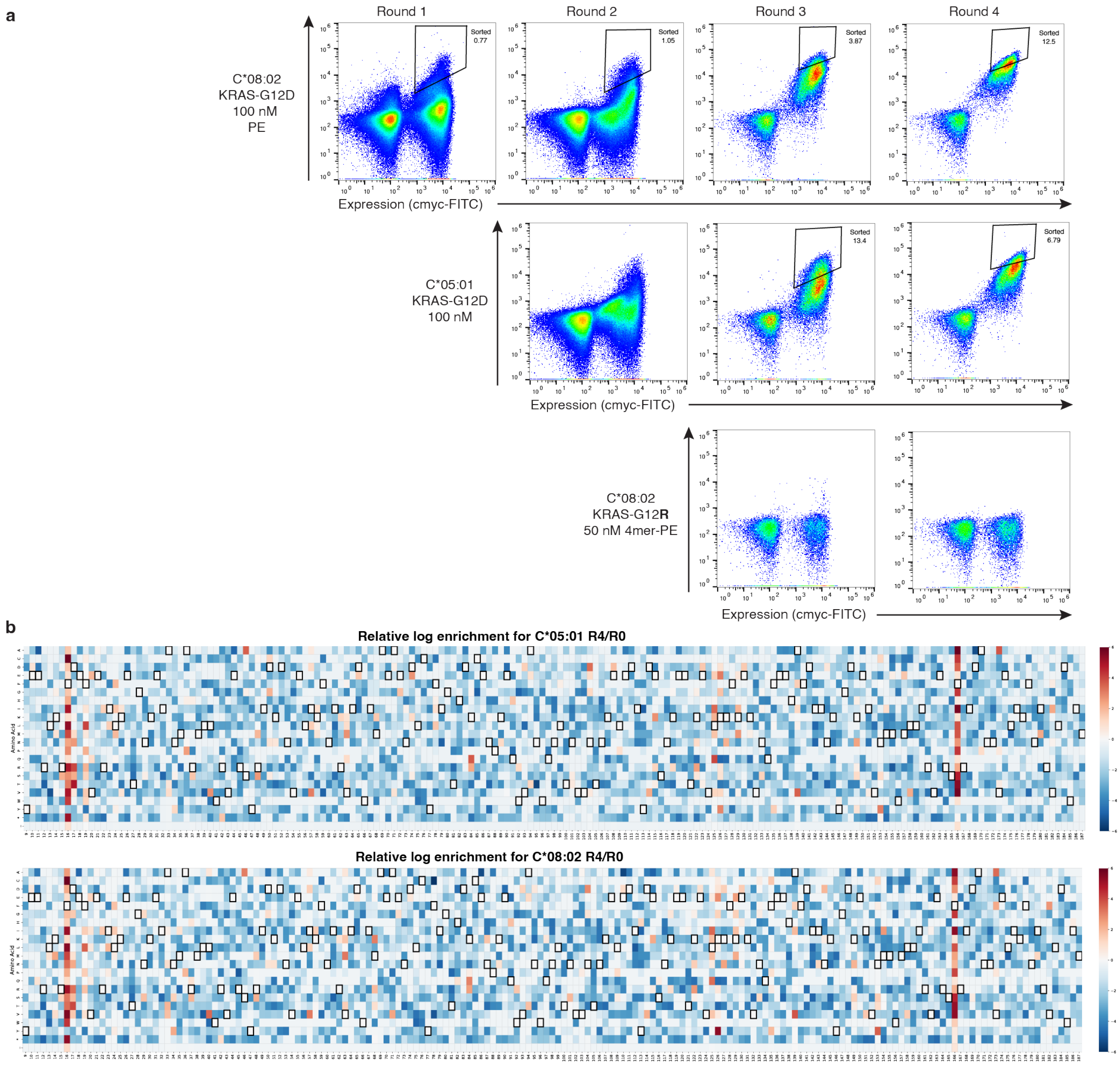


**Extended Data Fig. 10. a. Development of xTR-KRAS$^{HA}$ using error-prone mutagenesis.** Monovalent pHLA targets were used to select for high-affinity variants, removing reliance on avidity. **b.** Per-residue log enrichment from round 4 library versus round 0 library quantified using next-generation sequencing. Two positions F16 and F166 are frequently mutated to less bulky residues such as cysteine, valine, leucine, and isoleucine across the enriched sequences.

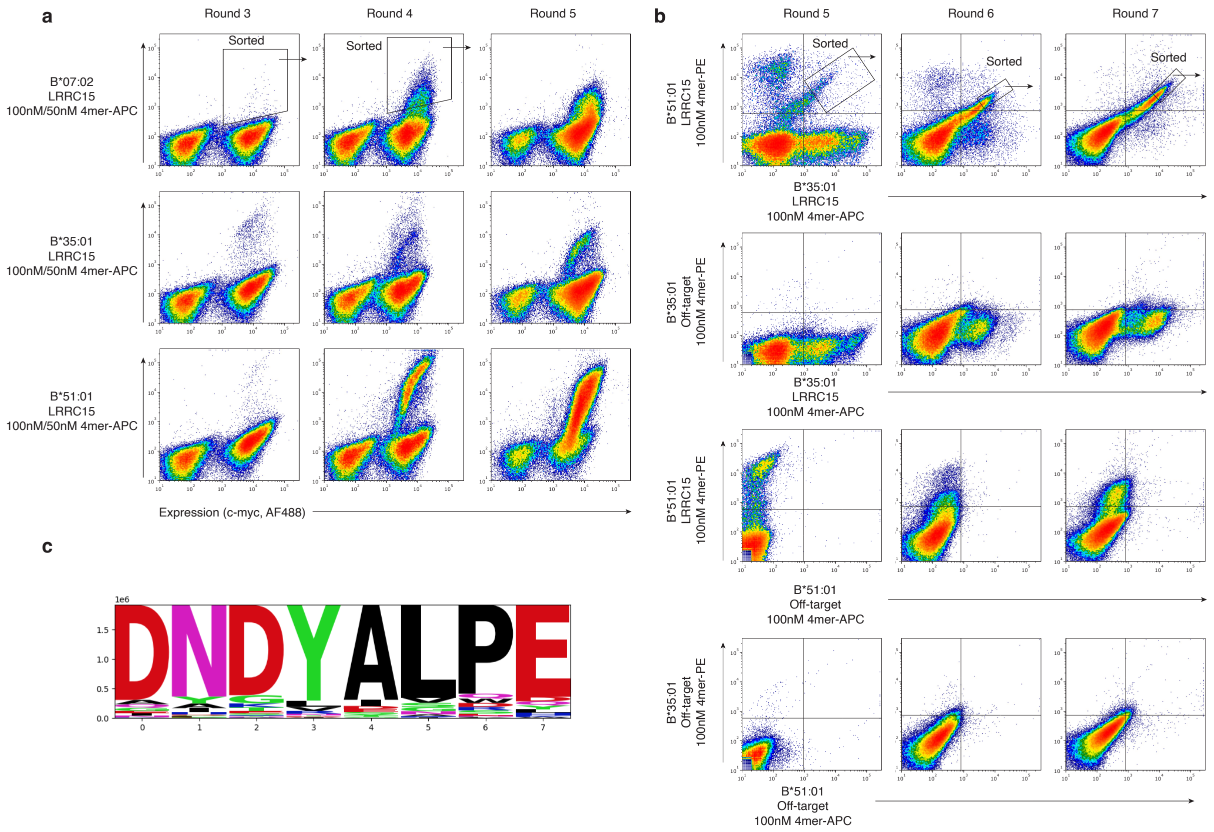


**Extended Data Fig. 11. Development of xTR-LRRC15. a.** Enrichment process from naïve TRACeR library targeting a peptide antigen from *LRRC15* (MPLKHYLLL) across HLA-B*07:02, HLA-B*35:01, and HLA-B*51:01 allotypes using single color sorting **b.** After round 4, dual-fluorophore staining in which equimolar ratios of orthogonal pHLA tetramers were incubated with yeast were sorted to isolate cross-HLA specific clones. **c.** Sequence analysis of top clones coming from round 7 double-positive population, with specificity to HLA-B*35:01/LRRC15 and HLA-B*51:01/LRRC15.

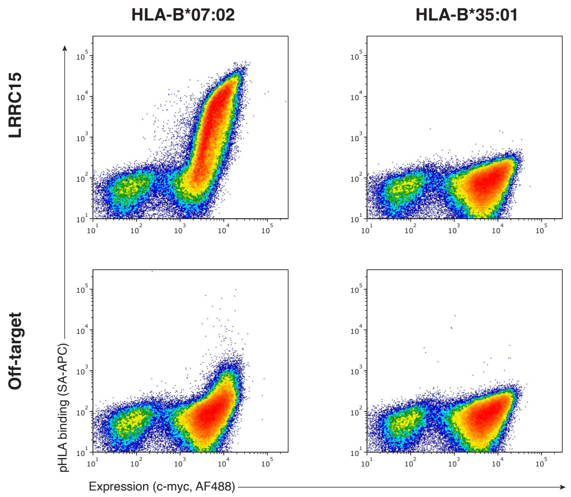


**Extended Data Fig. 12. Enrichment of specific binders for HLA-B*07:02/LRRC15.** Selections can isolate orthogonal yeast populations that specifically bind to HLA-B*07:02/LRRC15, but do not show cross-HLA binding to HLA-B*35:01/LRRC15. Here, LPNPIRTTA was used as an irrelevant, off-target peptide to demonstrate peptide specificity.

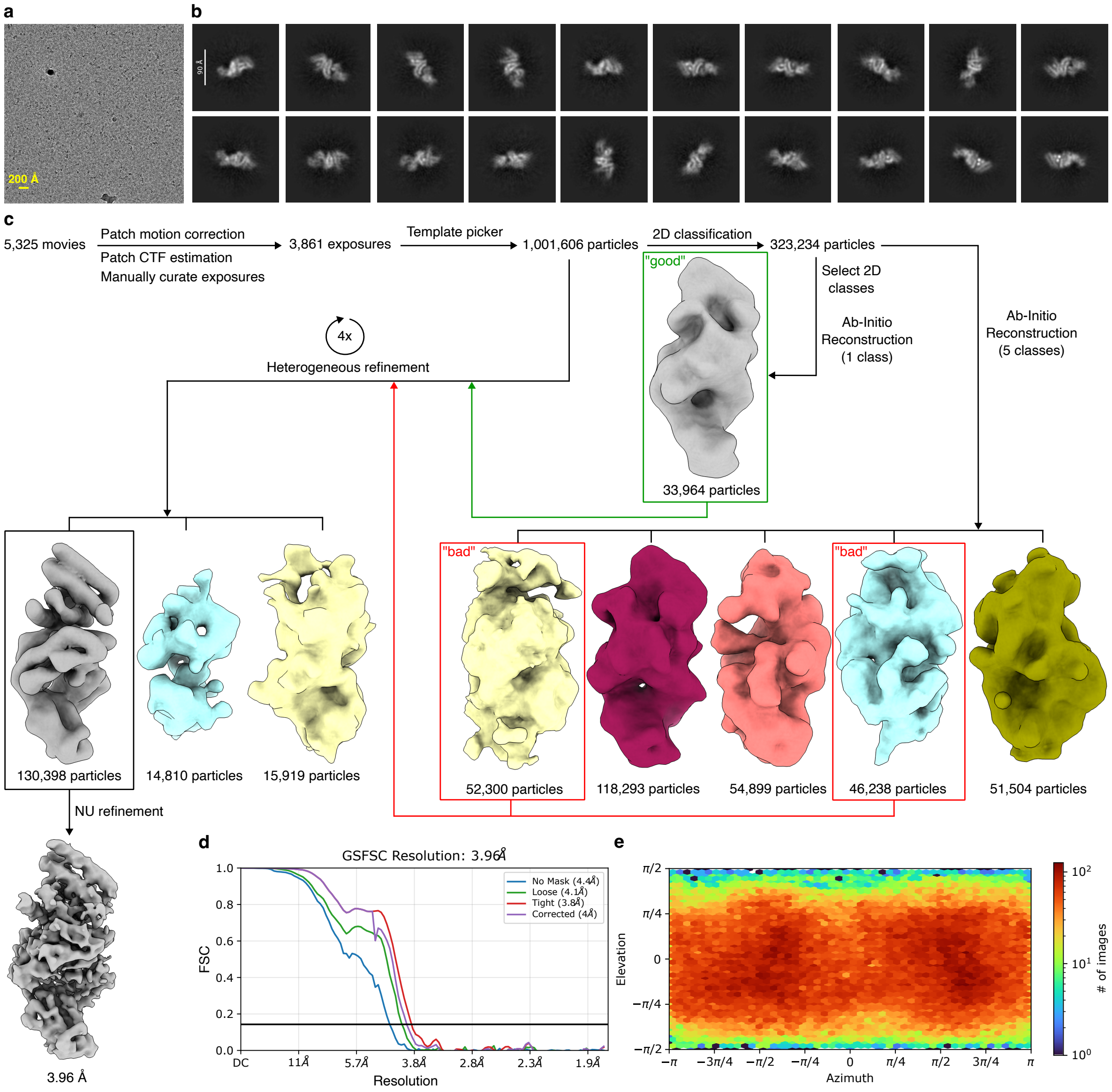


**Extended Data Fig. 13. Cryo-EM processing workflow for xTR-PRAME$^{HA}$/PRAME/HLA-A*11:01/β$_2$m complex. a.** Representative micrograph of the complex. **b.** 2D class averages of the complex particles. **c.** Data processing flowchart. **d.** Gold-standard Fourier shell correlation curves of the final reconstruction. **e.** Angular distribution plot.

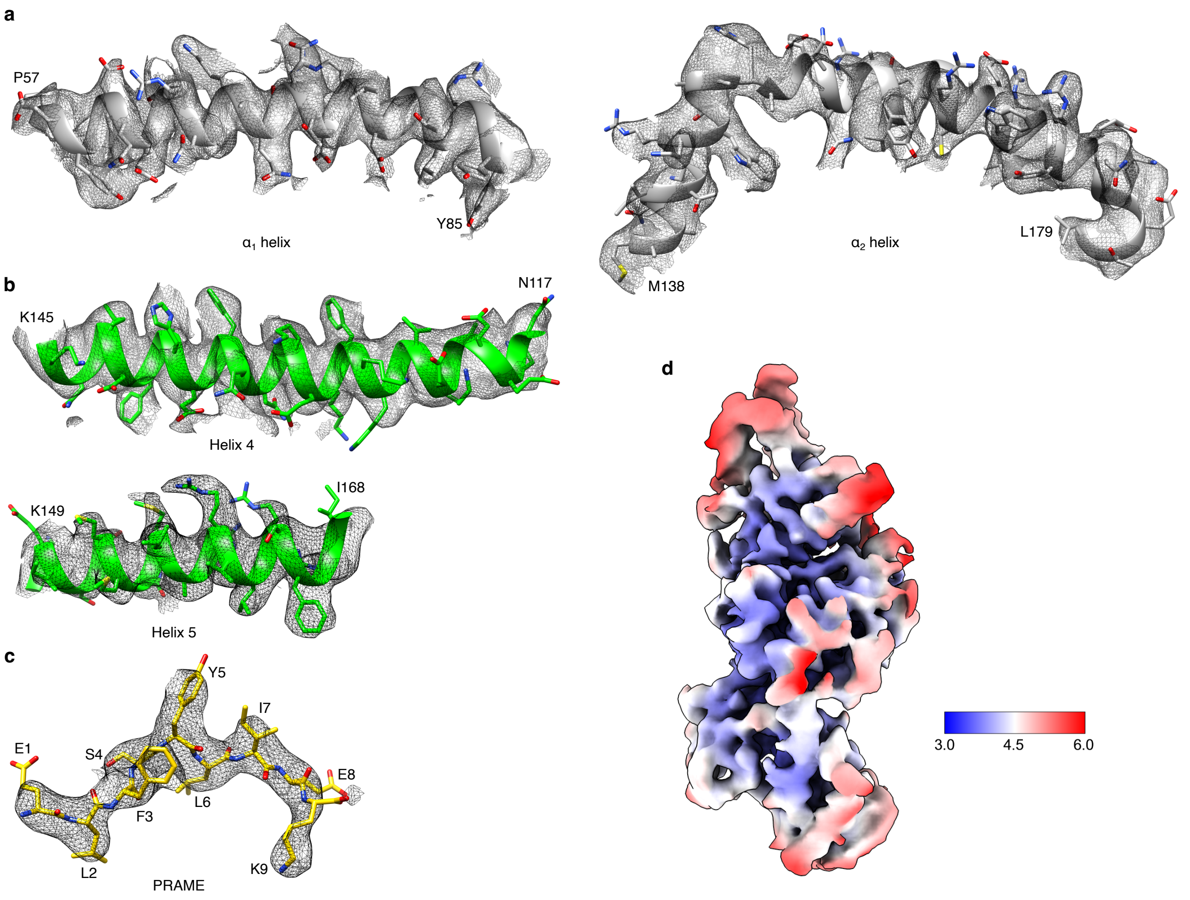


**Extended Data Fig. 14. Map and model quality for xTR-PRAME$^{HA}$/PRAME/HLA-A*11:01/β$_2$m complex. a.** Cryo-EM densities for $\alpha_1$ and $\alpha_2$ helices of HLA-A*11:01 at contour levels of 0.22 and 0.15, respectively. **b.** Cryo-EM densities for helices 4 and 5 of xTR-PRAME$^{HA}$ interacting with pHLA surface at contour levels of 0.14 and 0.17, respectively. **c.** Cryo-EM density of the PRAME epitope at contour level of 0.22. **d.** Local resolution distribution of the complex (Scale bar is represented in Å).

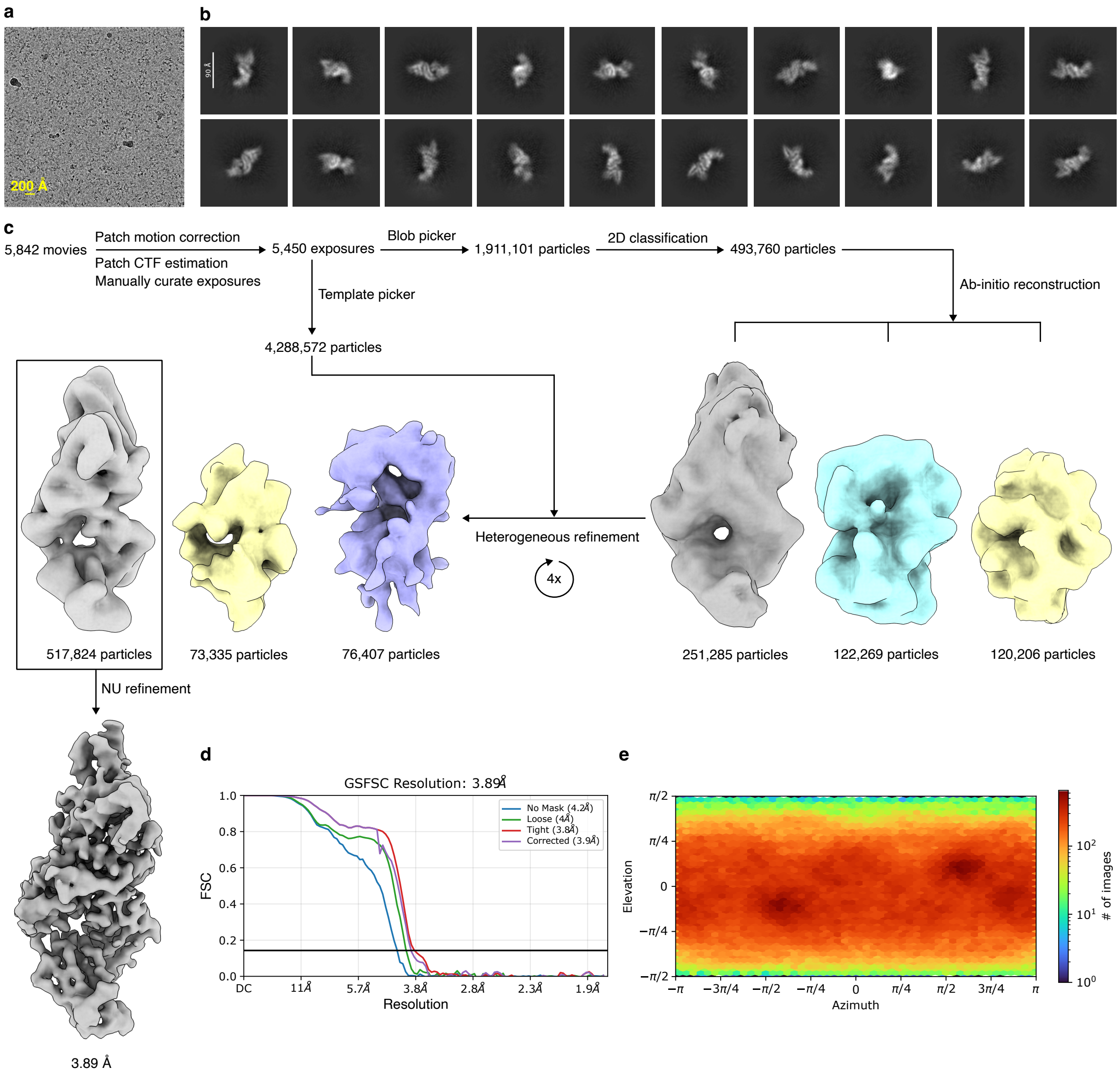


**Extended Data Fig. 15. Cryo-EM processing workflow for xTR-PHOX2B$^{HA}$/PHOX2B/HLA-C*07:02/β$_2$m complex. a.** Representative micrograph of the complex. **b.** 2D class averages of the complex particles. **c.** Data processing flowchart. **d.** Gold-standard Fourier shell correlation curves of the final reconstruction. **e.** Angular distribution plot.

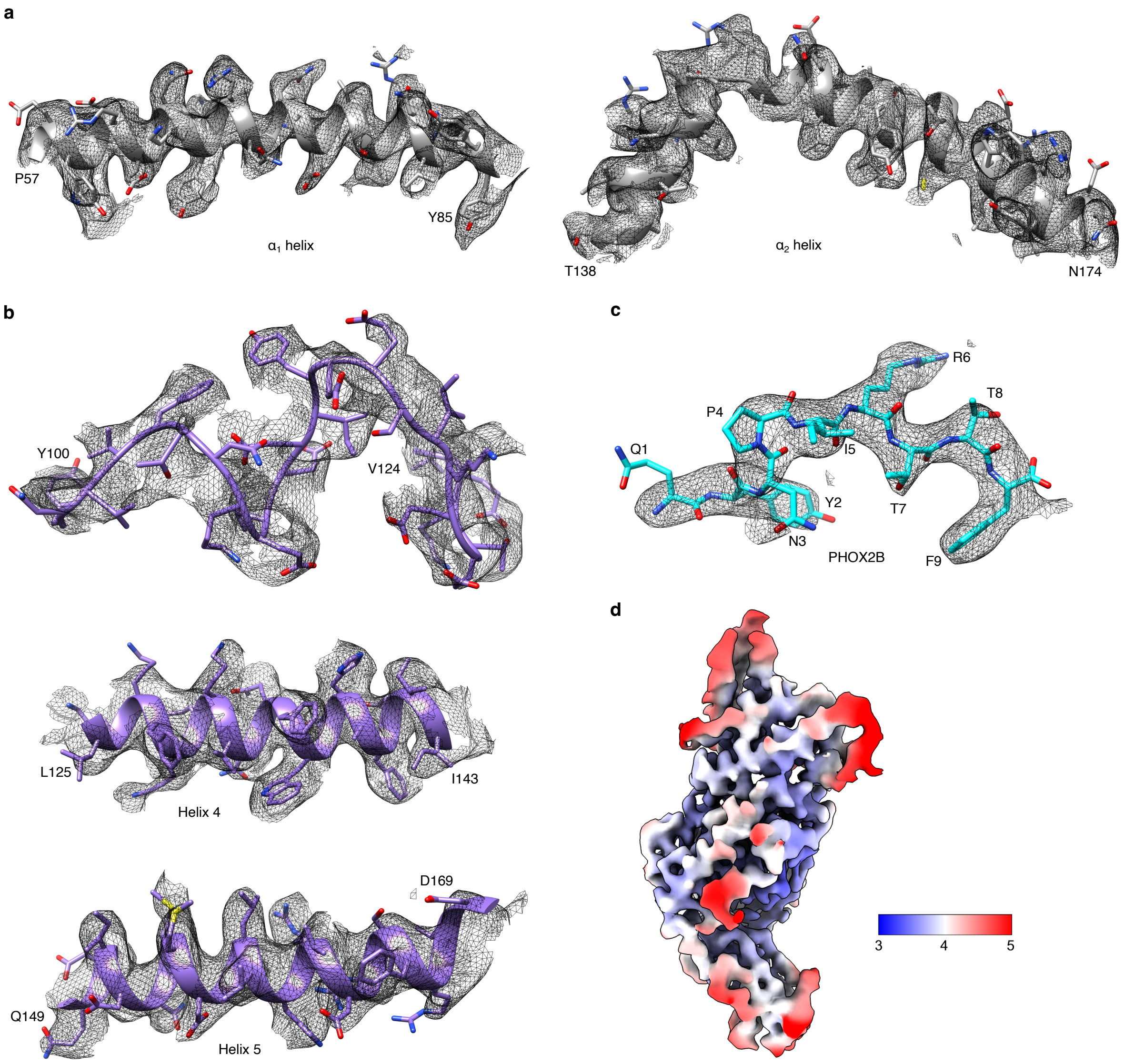


**Extended Data Fig. 16. Map and model quality for xTR-PHOX2B$^{HA}$/PHOX2B/HLA-C*07:02/$\beta_2$m complex. a.** Cryo-EM densities for $\alpha_1$ and $\alpha_2$ helices of HLA-C*07:02 at contour levels of 0.167 and 0.145, respectively. **b.** Cryo-EM densities for unstructured region (Y100-V124), helix 4, and helix 5 of xTR-PHOX2B$^{HA}$ interacting with pHLA surface at contour levels of 0.143, 0.120, and 0.161, respectively. **c.** Cryo-EM density of the PHOX2B epitope at contour level of 0.224. **d.** Local resolution distribution of the complex (Scale bar is represented in Å).

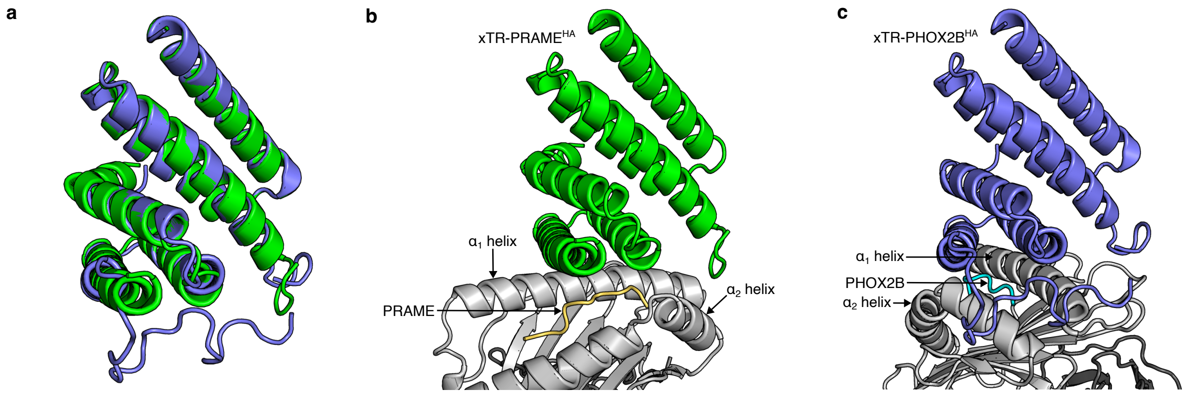


**Extended Data Fig. 17. xTR-PRAME$^{HA}$ and xTR-PHOX2B$^{HA}$ adopt same overall fold with different docking angles on their pHLA targets. a.** Structural superposition of xTR-PRAME$^{HA}$ (green) and xTR-PHOX2B$^{HA}$ (violet) showing that the overall helical bundle domain architecture of the TRACeR scaffold is maintained. **b.** Side-view of xTR-PRAME$^{HA}$ bound to HLA-A*11:01 presenting PRAME epitope. **c.** Side-view of xTR-PHOX2B$^{HA}$ bound to HLA-C*07:02 presenting PHOX2B epitope.

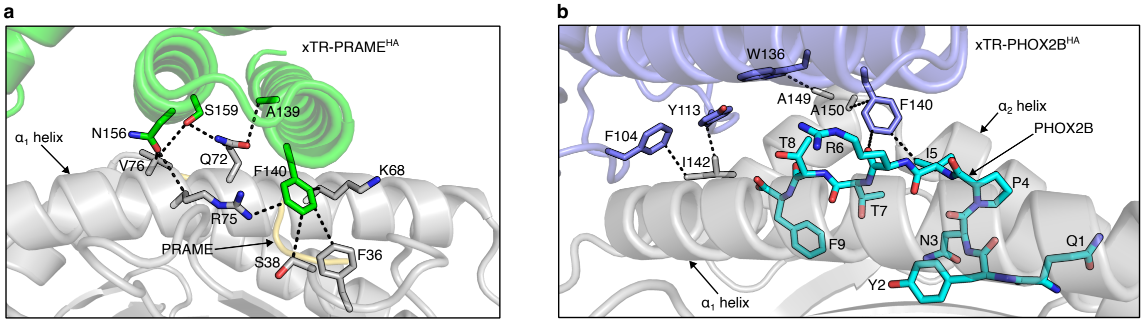


**Extended Data Fig. 18. Nonpolar interactions at the TRACeR/pHLA interface. a.** Nonpolar interactions mediated by xTR-PRAME$^{HA}$ with HLA-A*11:01 are shown in black dashed lines. **b.** Nonpolar interactions mediated by xTR-PHOX2B$^{HA}$ with HLA-C*07:02 and PHOX2B epitope are shown in black dashed lines.

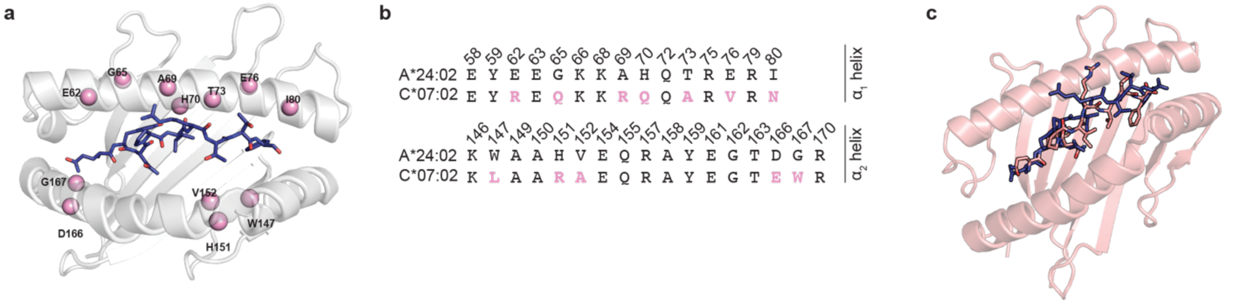


**Extended Data Fig. 19. Comparison of polymorphic surface residues and PHOX2B peptide conformations presented across HLA-A*24:02 and HLA-C*07:02. a.** Polymorphisms on framework residues spanning solvent-exposed positions along the $\alpha_1$ and $\alpha_2$ helices of HLA-A*24:02 and HLA-C*07:02. **b.** Sequence alignment of surface-exposed residues between HLA-A*24:02 and HLA-C*07:02, with polymorphisms shown as pink spheres in **(a)** outlined in pink. **c.** Overlay of PHOX2B peptide conformation on PHOX2B/HLA-A*24:02 complex (violet, PDB: 8EK) and PHOX2B/HLA-C*07:02 complex (salmon, PDB: 13FH). The high structural divergence of the peptide conformation between the two complexes is highlighted by a D-score of 5.97.

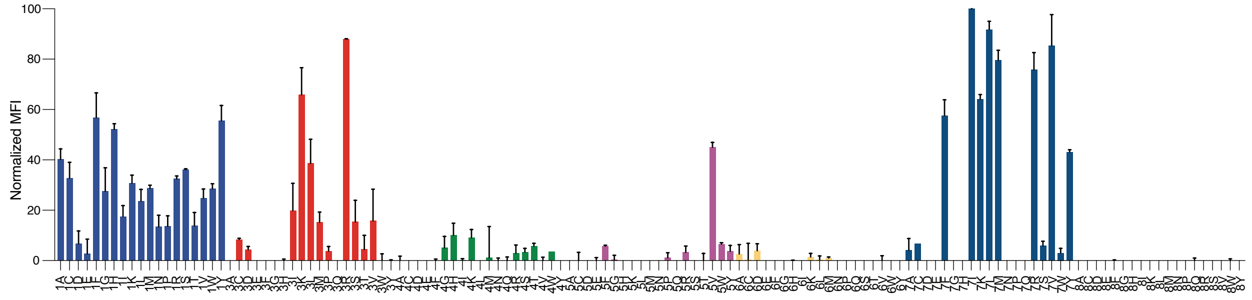


**Extended Data Fig. 20. xTR-PHOX2B$^{HA}$ peptide cross-reactivity analysis.** X-scan of non-anchor PHOX2B peptide residues (QYNPIRTTF, excluding Y2 and F9), using HLA-C*07:02 monoallelic B721.221 cells pulsed with 10 μM of each indicated PHOX2B point mutant, and stained with tetramerized, fluorescently labeled xTR-PHOX2B$^{HA}$. Bar chart displays binding MFI normalized to the highest value, adjusted for background staining. Data is shown as mean ± SD of n=2 technical replicates.

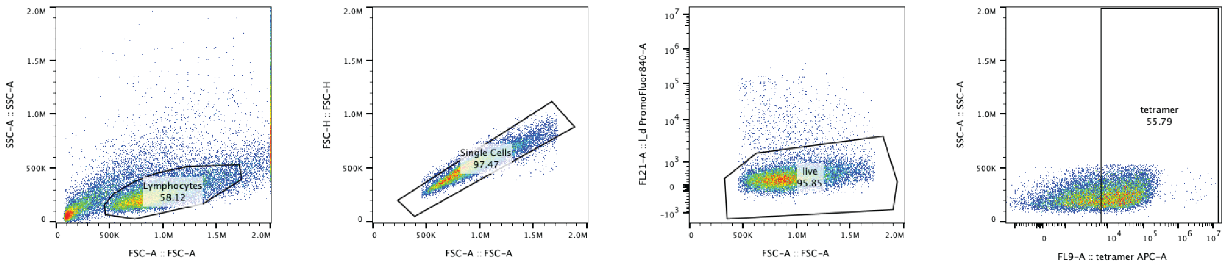


**Extended Data Fig. 21. Gating strategy for TRACeR-CAR positive primary T cells or $J^{ASP90}$ Jurkat reporter cells using on-target pHLA tetramers.** Starting event populations were gated under FSC-A/SSC-A plots for cells. T cell singlets were identified using FSC-A/FSC-H plots. Live cells were then gated using FSC-A/Live-dead Near IR plots. Tetramer positive cells were finally gated using Tetramer APC-A/SSC-A plots.

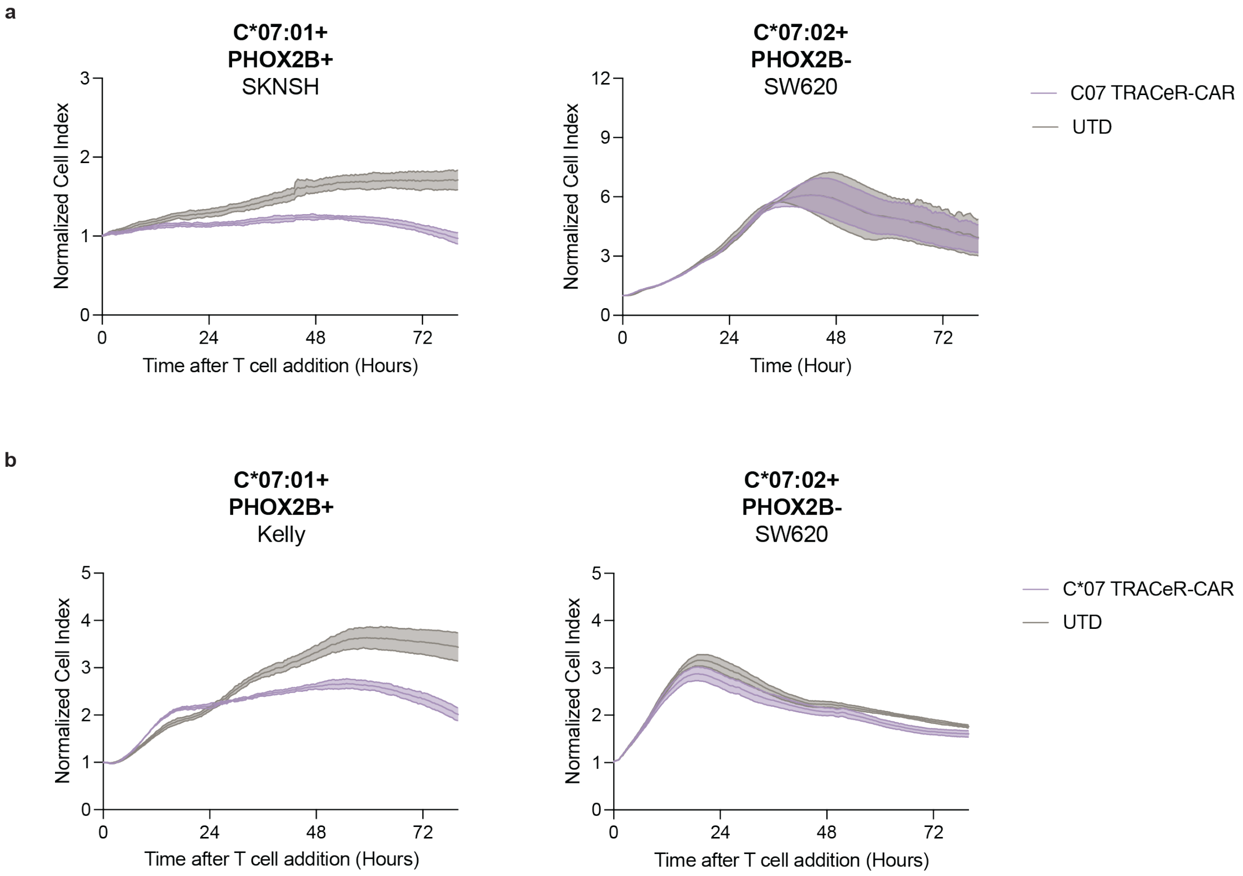


**Extended Data Fig. 22. Independent donor results for *in vitro* xTR-PHOX2B$^{HA}$-CAR T cell killing. a.** Cytotoxicity of xTR-PHOX2B$^{ha}$-CAR T cells co-cultured at 1:1 E:T with an HLA-C*07:01 expressing neuroblastoma cell line (SKNSH) and a PHOX2B-negative off-target cell line, measured by impedance. Data are shown as mean ± SD of 4 technical replicates. **b.** Same as (a) but for another HLA-C*07:01 expressing neuroblastoma cell line (Kelly). Data are shown as mean ± SD of 4 technical replicates.

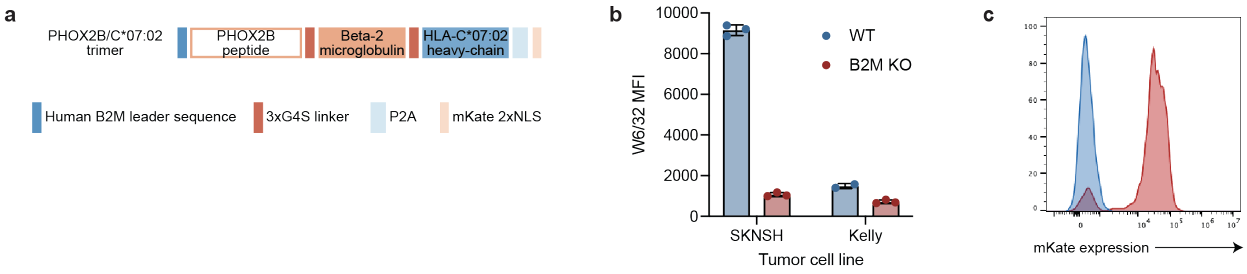


**Extended Data Fig. 23. HLA expression levels and pHLA enhancement strategy in established tumor cell lines. a.** Design of single-chain PHOX2B-$\beta_2$m-HLA-C*07:02 trimer. **b.** Anti-W6/32 stain of tumor cell lines with and without beta-2 microglobulin CRISPR knockout to validate abrogation of HLA expression. **c.** mKate expression to test for successful PHOX2B-$\beta_2$m-HLA-C*07:02 trimer transduction in SK-N-SH cells.

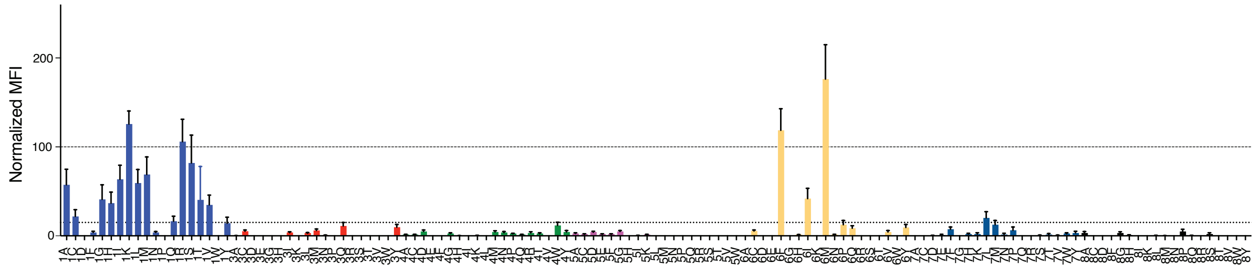


**Extended Data Fig. 24. xTR-PRAME$^{HA}$ peptide cross-reactivity analysis**. X-scan of non-anchor PRAME peptide residues, using HLA-A*11:01 phycoerythrin (PE) tetramers loaded with the indicated point mutant peptides (relative to ELFSYLIEK, excluding L2 and K9) to stain a xTR-PRAME$^{HA}$ expressing yeast clone. Bar chart displays binding MFI relative to HLA-A*11:01 loaded with WT PRAME peptide (set to 100%), adjusted for background staining. Dashed line indicates the 15% MFI cutoff used to define a regular expression for searching the human proteome for potentially cross-reactive self-peptides. This cutoff yielded the motif [ACGHIKMPQRSTVYE]-[VTISLA]-[QYF]-[WS]-Y-[EFIMYL]-[CLM]-E-[KRY], resulting in one potential off-target peptide, MLFWYIMEK encoded by the *MROH2B* human gene. Data is shown as mean ± SD of n=3 technical replicates.

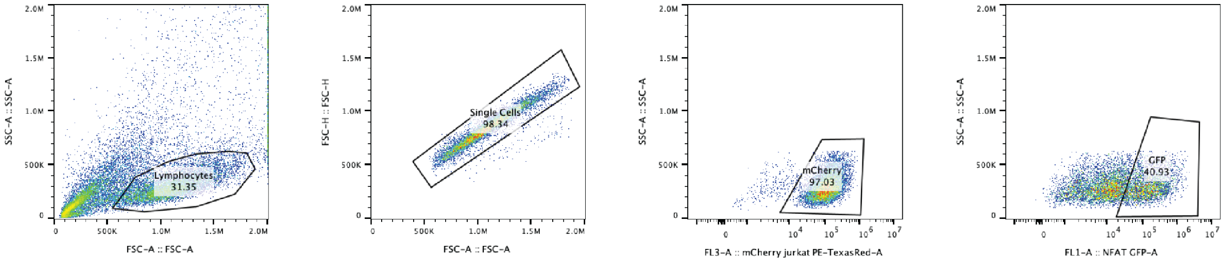


**Extended Data Fig. 25. J$^{ASP90}$ Jurkat reporter cell activation assay gating strategy.** Starting event populations were gated under FSC-A/SSC-A plots for cells. T cell singlets were identified using FSC-A/FSC-H plots. Jurkat cells were further identified using mCherry (PE-TexasRed)/SSC-A plots. Activated Jurkat cells were finally gated using GFP-A/SSC-A plots. GFP gate was set using unstimulated Jurkat cells and PMA-I stimulated Jurkat cells as negative and positive controls, respectively.

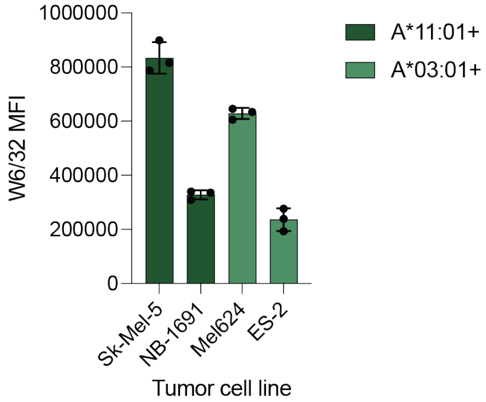


**Extended Data Fig. 26. HLA expression levels for cancer cell lines used in xTR-PRAME$^{HA}$-CAR T cell studies.** Anti-W6/32 stain of PRAME-expressing tumor cell lines shows variable levels of HLA expression.

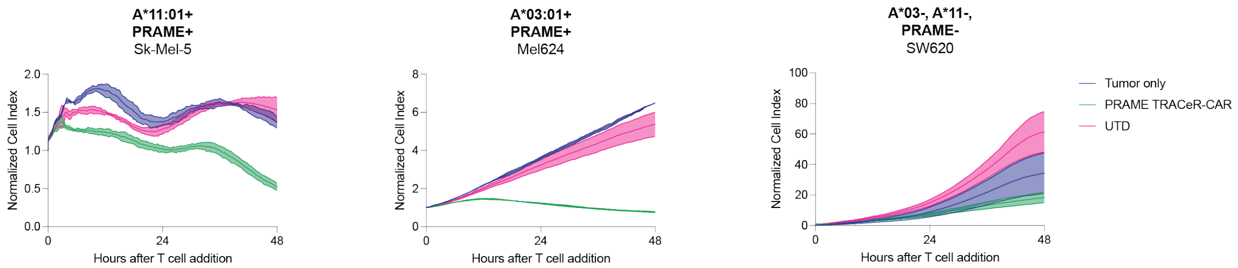


**Extended Data Fig. 27. Independent donor results for *in vitro* xTR-PRAME$^{HA}$-CAR T cell killing.** Cytotoxicity of xTR-PRAME$^{ha}$-CAR T cells co-cultured at 1:2 E:T with PRAME-positive melanoma cell lines and a PRAME-negative off-target cell line, measured by impedance. Data are shown as mean ± SD of 4 technical replicates.

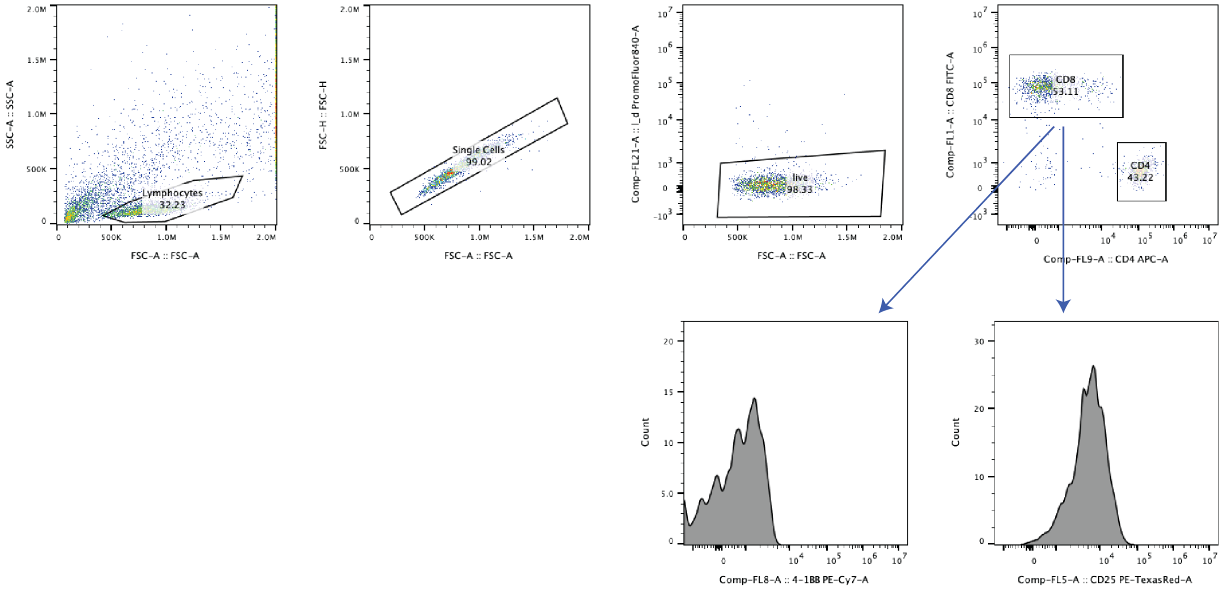


**Extended Data Fig. 28. Gating strategy for xTR-PRAME$^{HA}$-CAR T cell activation markers.** Starting event populations were gated under FSC-A/SSC-A plots for cells. T cell singlets were identified using FSC-A/FSC-H plots. Live cells were then gated using FSC-A/Live-dead Near IR plots. T cells were further gated on CD4 APC-A/CD8 FITC-A. MFI of each activation marker (CD25/4-1BB) was determined within CD4+ and CD8+ populations separately.

**Extended Data Table 1. Isothermal titration calorimetry parameters for PHOX2B.**

| | Replicate 1 | Replicate 2 |
|---|---|---|
| **$K_D$ (nM)** | 115.7 | 201.21 |
| **ΔH (cal/mol)** | -5523 ±254.8 | -5372 ±414.0 |
| **ΔS (cal/mol*deg)** | 13.2 | 12.6 |

**Extended Data Table 2. DSF melting temperatures for recombinant pHLA reagents used.**

| pHLA | Melting Temperature ($^{o}$C) |
|---|---|
| HLA-A*03:01/PRAME | 44.17 |
| HLA-A*11:01/PRAME | 52.68 |
| HLA-A*68:01/PRAME | 55.00 |
| HLA-A*03:01/VVGAGGVGK | 46.43 |
| HLA-A*11:01/VVGADGVGK | 55.29 |
| HLA-A*68:01/AIFQSSMTK | 42.48 |
| HLA-A*23:01/PHOX2B | 70.70 |
| HLA-A*24:02/PHOX2B | 63.20 |
| HLA-B*07:02/LRRC15 | 60.40 |
| HLA-B*35:01/LRRC15 | 58.22 |
| HLA-B*51:01/LRRC15 | 56.98 |
| HLA-B*07:02/ GADGVGKSAL | 59.34 |
| HLA-B*35:01/LPNPIRTTA | 51.18 |
| HLA-B*51:01/LPNPIRTTA | 55.27 |
| HLA-C*07:01/PHOX2B | 45.26 |
| HLA-C*07:02/PHOX2B | 45.53 |
| HLA-C*07:01/PHOX2B_R6A | 53.39 |
| HLA-C*07:02/PHOX2B_R6A | 52.12 |
| HLA-C*05:01/KRAS G12D 10mer | 40.27 |
| HLA-C*08:02/KRAS G12D 10mer | 52.10 |
| HLA-C*05:01/ELDEISTNI | 47.35 |
| HLA-C*08:02/GAGGVGKSAL | 58.57 |

### Appendix

#### PepPred Development and Training

Backbone dihedral angles of all peptide residues extracted from Protpardelle/[1]AFF2[2–4] models were projected onto 3D space via principal component analysis. Analyses were done independently for each group of HLA allotypes presenting a shared peptide antigen to capture dominant ensemble variation. For each projection, centroid position ($C_E$), the mean and trace of the spread of each ensemble ($E$) within the 3D space were calculated from individual members ($z_i$) using equations 1-4 below.

$$E = \{z(a,b,c)_i \dots z(a,b,c)_n\} \tag{1}$$

$$C_E = \frac{1}{n}\sum_{i=1}^{n} z(a,b,c)_i \tag{2}$$

$$spread_{mean}(E) = \frac{1}{n}\sum_{i=1}^{n} ||z(a,b,c)_i - c_E|| \tag{3}$$

$$spread_{trace}(E) = \sum_{i=1}^{n} Var(z(a,b,c)_i) \tag{4}$$

$$D(\theta_1, \theta_2) = 2(1 - \cos(\theta_1 - \theta_2)) \tag{5}$$

$$D - score(A,B) = \sum_{p=4}^{7} D(\phi_p^A, \phi_p^B) + D(\psi_p^A, \psi_p^A) \tag{6}$$

Pairwise D-Scores [3,5](equations 5-6) computed for each 104 pHLA pair were used to establish a ground truth set, whereby pairs with scores less than 1.5 were classified as similar in terms of their overall peptide conformation and scores of 1.5 or greater were classified as dissimilar during training based on our previous benchmark[3]. For each paired pHLA ensemble, pairwise comparative features were generated from the geometric descriptors of the ensembles in the low-dimensional space (Equations 1-4), as well as confidence metrics from the initial AFF2 structural predictions (iPTM, PAE matrices, pLDDT scores) (**Appendix Fig. 1**). Additional aggregate dihedral information was also passed to training via calculated minimums, maximums, and mean values of $\phi$ and $\psi$ dihedral angles for each residue of the peptide chain for each ensemble. The ground truth set of 104 labeled peptide/HLA pairs was used to train a stacked ensemble of Elastic Net logistic regression[6], Gradient Boosting[7], and Gaussian Process classifiers[8], which produced outputs that were interpreted by a Ridge Regression meta-learner[9] to yield *predicted peptide similarity scores* for each pHLA pair in the training set (**Appendix Fig. 2**). The final meta learner additionally formed classifications per pHLA pair utilizing optimized threshold values to maximize recovery and label accuracy. The model training and evaluation was done using 5-fold cross validation, context-aware leave one out cross validation, and label-shuffling to prevent data leakage and assess accuracy to unseen structural contexts (**Appendix Fig. 3**). Leave one out cross validation was made context-aware by grouping pairs based on shared peptide antigen to ensure consistency on unseen antigen contexts. During execution of the method for a new peptide target with confirmed or predicted presentation across a set of HLA allotypes, the AFF2/Protpardelle structural ensemble generation process is first run for each pHLA in the set using a distributed computing system. PepPred then evaluates predicted peptide similarity scores for each pHLA pair, and results are plotted as heatmaps. Alleles on heatmap are clustered iteratively by grouping pairs of the most similar clusters and alleles to form a dendrogram.

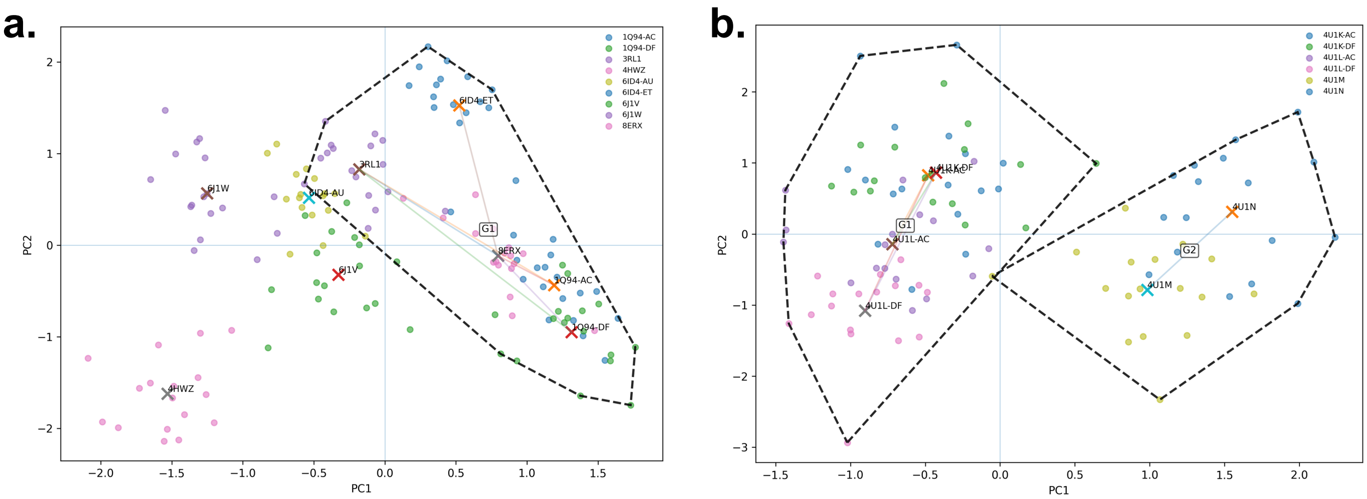


**Appendix Fig. 1. Feature extraction from Principal Component Analysis of pHLA Structural Ensembles. a.** A two-dimensional visualization of PCA performed for all Protpardelle/AFF2 predicted structural ensembles for alleles presenting peptide AIFQSSMTK. Ensembles are labelled by corresponding solved crystal structures which serve as ground truth representatives during training. Centroid positions for each ensemble are presented as cross-marks, and centroid distances of ground-truth labeled similar ensembles are shown as linear connections between centroids. Spread of ensembles corresponding ground-truth determined similar structures is outlined along shared edges to define "G1" cluster of similarly conformed ensembles. Spread of individual ensembles, centroid positions, and centroid distances between pairs are passed onto an L2 Elastic Net learner for training. **b.** Two-dimensional visualization and annotation of PCA performed for all Protpardelle/AFF2 predicted structural ensembles for alleles presenting peptide RPQVPLRPM. Two clusters are defined by grouping ensembles by shared edges along ground-truth determined similarity of corresponding solved crystal structures.

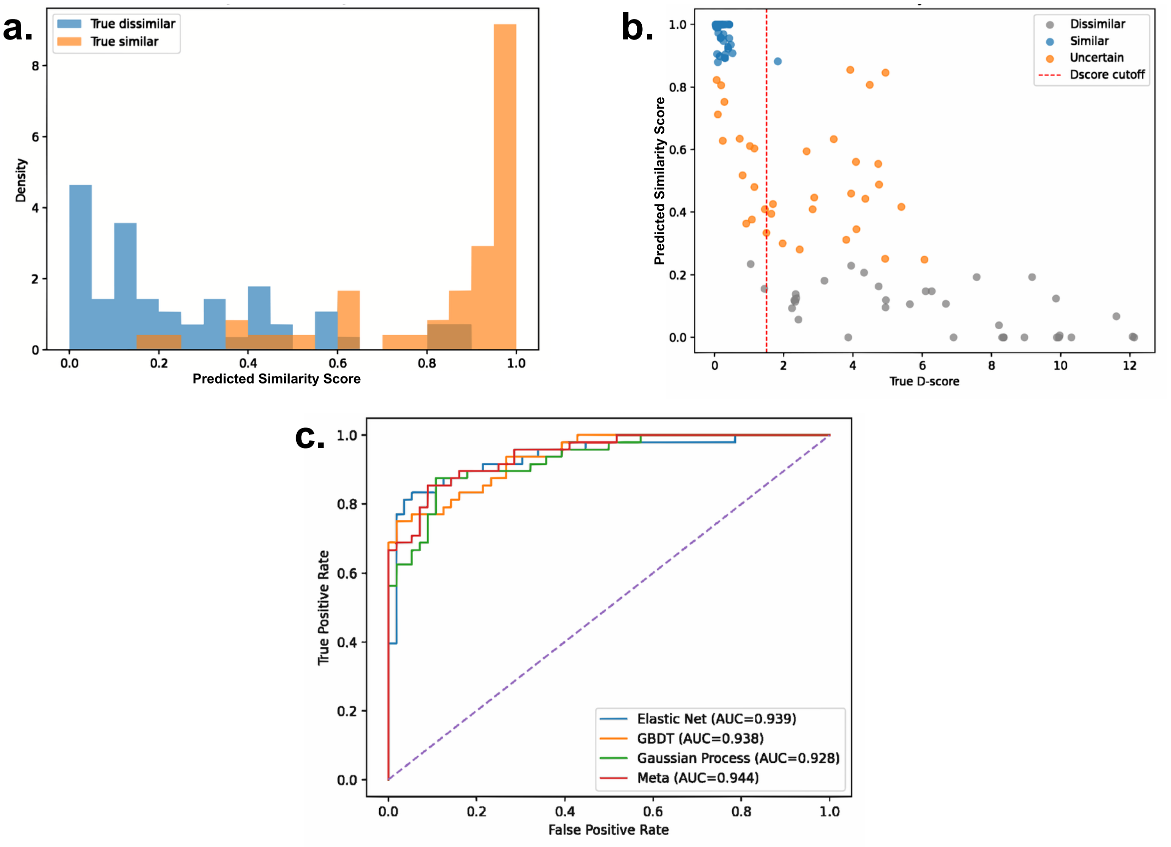


**Appendix Fig. 2. PepPred Model Performance and Accuracy. a.** Predicted Similarity Scores for pHLA pairs calculated by the Ridge Regression meta-learner were plotted by ground truth classification. Densities for pairs classified as “True Similar” based on D-Score of corresponding ground truth structures exhibit greatest density at scores above 0.8, whereas pairs classified as “True Dissimilar” exhibit greatest densities below scores of 0.2. **b.** Predicted Similarity Scores for pHLA pairs calculated by Ridge Regression meta-learner were plotted against ground truth D-Scores and classified into three optimized buckets of model confidence. Threshold values for classification of pairs were optimized to preserve model accuracy in “Similar” and “Dissimilar” buckets, creating an “Uncertain” class for pairs with low confidence separation. Cuttoffs were further optimized to maximize recovery yielding a single definitional threshold and generate a balanced cutoff used to produce the final “Similar” and “Dissimilar” classifications. **c.** ROC curve for each meta learner was plotted from final testing set for each base learner as well as Ridge Regression meta learner. Performance was measured on final test set classifications of the ensemble model after validation.

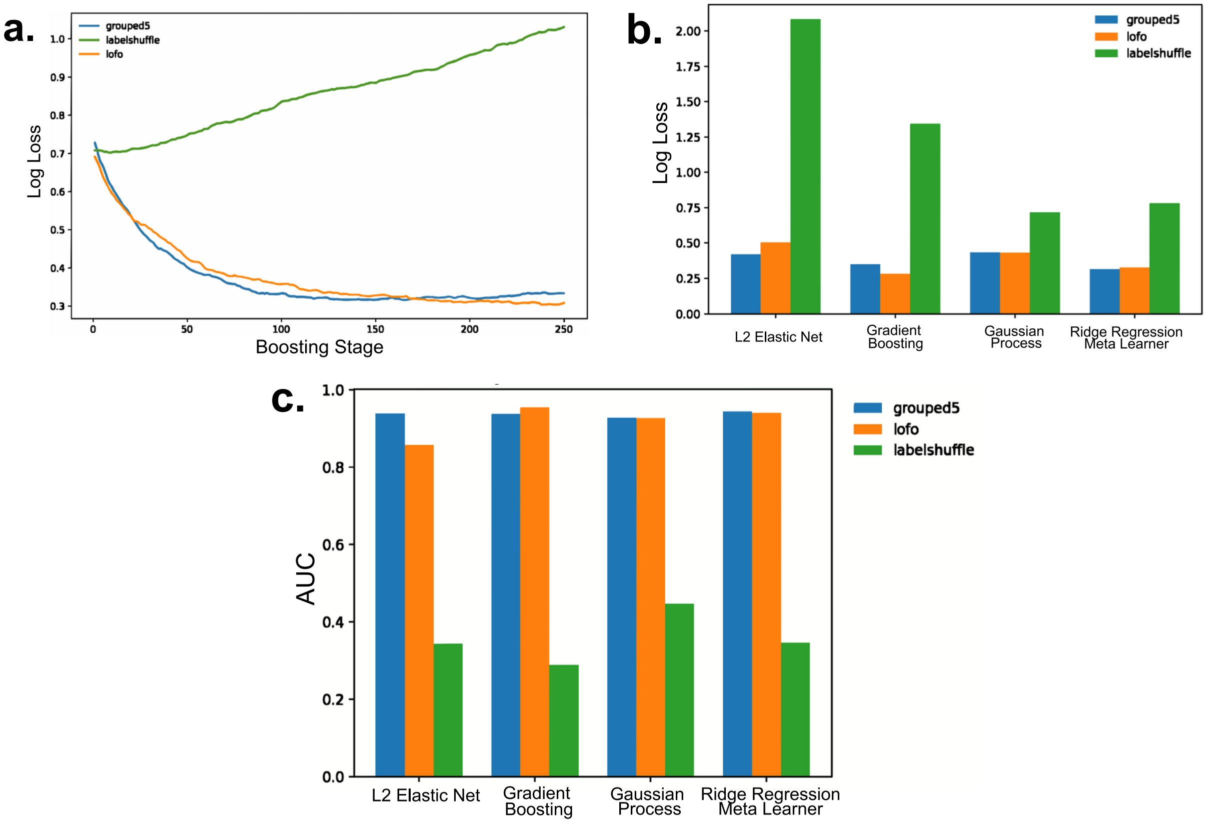


**Appendix Fig. 3. PepPred Model Validation. a.** Mean Log Loss for Out of Fold predictions at each boosting stage for Gradient Boosted Decision Tree base learner under three validation modes: 5-Fold Cross Validation (grouped5), Context-Aware Leave one Out Cross Validation (lofo), and Label Shuffling Validation (labelshuffle). Log Loss was calculated using binary cross-entropy (Equation 7) for out-of-fold predictions at every boosting stag. **b.** Mean Log Loss calculated at the best epoch or boosting stage for each component under 3 validation modes. **c.** Component model performance under validation modes given by mean AUC values calculated from ROC curves of model predictive accuracy in Out of Fold rounds.

$$Log\ Loss = -\frac{1}{N}\sum_{i=1}^{N}[y_i \log(p_i) + (1 - y_i)\log(1 - p_i)] \quad (7)$$
